# Sulfur-oxidizing symbionts without canonical genes for autotrophic CO_2_ fixation

**DOI:** 10.1101/540435

**Authors:** Brandon K. B. Seah, Chakkiath Paul Antony, Bruno Huettel, Jan Zarzycki, Lennart Schada von Borzyskowski, Tobias J. Erb, Angela Kouris, Manuel Kleiner, Manuel Liebeke, Nicole Dubilier, Harald R. Gruber-Vodicka

**Affiliations:** Max Planck Institute for Marine Microbiology, Celsiusstraße 1, 28359 Bremen, Germany; Max Planck Genome Centre Cologne, Max Planck Institute for Plant Breeding Research, Carl-von-Linné-Weg 10, 50829 Cologne, Germany; Max Planck Institute for Terrestrial Microbiology, Karl-von-Frisch-Str. 10, 35043 Marburg, Germany; Energy Bioengineering and Geomicrobiology Group, University of Calgary, 2500 University Drive Northwest, Calgary, Alberta T2N 1N4, Canada; Department of Plant and Microbial Biology, North Carolina State University, Raleigh 27695, North Carolina, United States of America; MARUM, Center for Marine Environmental Sciences, University of Bremen, 28359 Bremen, Germany; Max Planck Institute for Developmental Biology, Max-Planck-Ring 5, 72076 Tübingen, Germany; Red Sea Research Center, Biological and Environmental Sciences and Engineering (BESE) Division, King Abdullah University of Science and Technology (KAUST), Thuwal 23955, Kingdom of Saudi Arabia

**Keywords:** meiofauna, ectosymbiont, Gammaproteobacteria, protist, lithoheterotrophy

## Abstract

Since the discovery of symbioses between sulfur-oxidizing (thiotrophic) bacteria and invertebrates at hydrothermal vents over 40 years ago, it has been assumed that autotrophic fixation of CO_2_ by the symbionts drives these nutritional associations. In this study, we investigated *Candidatus* Kentron, the clade of symbionts hosted by *Kentrophoros*, a diverse genus of ciliates which are found in marine coastal sediments around the world. Despite being the main food source for their hosts, Kentron lack the key canonical genes for any of the known pathways for autotrophic fixation, and have a carbon stable isotope fingerprint unlike other thiotrophic symbionts from similar habitats. Our genomic and transcriptomic analyses instead found metabolic features consistent with growth on organic carbon, especially organic and amino acids, for which they have abundant uptake transporters. All known thiotrophic symbionts have converged on using reduced sulfur to generate energy lithotrophically, but they are diverse in their carbon sources. Some clades are obligate autotrophs, while many are mixotrophs that can supplement autotrophic carbon fixation with heterotrophic capabilities similar to those in Kentron. We have shown that Kentron are the only thiotrophic symbionts that appear to be entirely heterotrophic, unlike all other thiotrophic symbionts studied to date, which possess either the Calvin-Benson-Bassham or reverse tricarboxylic acid cycles for autotrophy.

**Significance Statement:** Many animals and protists depend on symbiotic sulfur-oxidizing bacteria as their main food source. These bacteria use energy from oxidizing inorganic sulfur compounds to make biomass autotrophically from CO_2_, serving as primary producers for their hosts. Here we describe apparently non-autotrophic sulfur symbionts called Kentron, associated with marine ciliates. They lack genes for known autotrophic pathways, and have a carbon stable isotope fingerprint heavier than other symbionts from similar habitats. Instead they have the potential to oxidize sulfur to fuel the uptake of organic compounds for heterotrophic growth, a metabolic mode called chemolithoheterotrophy that is not found in other symbioses. Although several symbionts have heterotrophic features to supplement primary production, in Kentron they appear to supplant it entirely.

## Introduction

Chemosynthetic symbioses between heterotrophic, eukaryotic hosts and bacteria that use the oxidation of inorganic chemicals or methane to fuel growth are common in marine environments. They occur in habitats ranging from deep sea vents and seeps, where they are responsible for much of the primary production, to the shallow water interstitial, where the hosts are often small and inconspicuous meiofauna. Among the energy sources for chemosynthesis are reduced sulfur species like sulfide and thiosulfate, and such sulfur-oxidizing (thiotrophic) symbioses have convergently evolved multiple times (1). They are commonly interpreted as nutritional symbioses where the symbionts fix CO_2_ autotrophically into biomass with the energy from sulfur oxidation and eventually serve as food for their hosts (1, 2). Indeed, several host groups have become so completely dependent on their symbionts for nutrition that they have reduced digestive systems. All sulfur-oxidizing symbioses investigated thus far possess a primary thiotrophic symbiont with genes of either the Calvin-Benson-Bassham (CBB) (3–10) or reverse tricarboxylic acid (rTCA) (11, 12) cycles for CO_2_ fixation, and the different pathways may relate to different ecological niches occupied by the symbioses (13). The symbionts of the vestimentiferan tubeworms are additionally able to encode both the CBB and rTCA cycles, which may be active under different environmental conditions (14–16). Beyond sulfur oxidation and carbon fixation, several thiotrophic symbionts have additional metabolic capabilities such as the uptake of organic carbon (17), the use of carbon monoxide (18) and hydrogen (19) as energy sources, and the ability to fix inorganic nitrogen (4, 5).

The thiotrophic ectosymbionts of the ciliate genus *Kentrophoros* constitute a distinct clade of Gammaproteobacteria named “*Candidatus* Kentron” (hereafter Kentron) (20). Kentron has previously been shown to oxidize sulfide and fix CO_2_ (21), and to be consumed and digested by its hosts (21, 22). Unlike most ciliates, which consume their food at a specific location on the cell that bears feeding structures composed of specialized cilia, *Kentrophoros* has only vestiges of such cilia, and instead directly engulfs its symbionts along the entire cell body (23), suggesting that Kentron bacteria are its main food source.

Given that all previous studies of thiotrophic symbionts, including Kentron, have characterized them as autotrophic, we expected that the pathways of energy and carbon metabolism used by Kentron would resemble those in other thiotrophic bacteria involved in nutritional symbioses. In this study, we used metagenomic and transcriptomic analyses of single host individuals to show that the Kentron clade lacks the canonical pathways of autotrophic CO_2_ fixation. Based on a metabolic reconstruction of the core genome from eleven Kentron phylotypes collected from three different sites, and results from direct protein stable isotope fingerprinting, we propose that it is a lithoheterotrophic nutritional symbiont, relying on assimilation of organic substrates rather than fixation of inorganic carbon to feed its hosts.

## Results

### Symbiont genome assemblies have high coverage and completeness, and represent eleven phylotypes

Genomes of Kentron symbionts were binned from 34 metagenome assemblies, each corresponding to a single *Kentrophoros* host ciliate individual. These samples represented 12 host morphospecies from three different geographical locations: the Mediterranean, Caribbean, and Baltic Seas (Supplementary Table 1). The symbiont genome assemblies had total lengths between 3.31 to 5.02 Mbp (median 3.91 Mbp), although they were relatively fragmented (N50: 3.52 to 37.5 kbp, median 21.4 kbp). Genome sizes and assembly fragmentation appeared to be species/phylotype-dependent (Supplementary Figure 1).

Nonetheless, the genome bins were relatively complete (91.4 to 94.9%, median 93.8%) and had low contamination (0.75 to 3.56%, median 1.87%) (Supplementary Table 2). The core genomic diversity in the clade was well-sampled: 1019 protein-coding gene orthologs were found in all 34 genomes, and the core genome accumulation curve reached a plateau (Supplementary Figure 2). Kentron genome sizes were relatively large for thiotrophic symbionts, and were comparable to values for *Ca*. Thiodiazotropha spp. (4.5 Mbp) and the Gamma3 symbiont of *Olavius algarvensis* (4.6 Mbp).

Kentron formed a well-supported clade (100% SH-like support value) within the Gammaproteobacteria, in a phylogenetic analysis using conserved protein-coding marker genes (Figure 1). Their closest relatives in the set of basal Gammaproteobacteria analysed were *Nitrosococcus oceani, Methylophaga thiooxydans, Thioploca ingrica, Ca*. Competibacter denitrificans, and *Beggiatoa* spp. (100% support), which differed from the 16S rRNA gene phylogeny, where Kentron was sister to the Coxiellaceae (20). Symbionts from different host morphospecies formed separate, well-supported phylotype clusters, with the exception of Kentron from *Kentrophoros* sp. UNK and *K*. sp. LPFa, where a single symbiont phylotype was associated with two different host phylotypes, as previously observed with 16S and 18S rRNA sequences. Among genomes of the same phylotype, average nucleotide identities (ANI) were 93.0–100% and average amino acid identities (AAI) were 93.2–100%, whereas between different phylotypes, these values were 83.2–93.8% and 70.6–91.3% respectively, which supports them being different species in the same genus (24). Kentron phylotypes will therefore be referred to here with their corresponding host morphospecies identifiers, except for Kentron UNK/LPFa.

**Figure 1.**
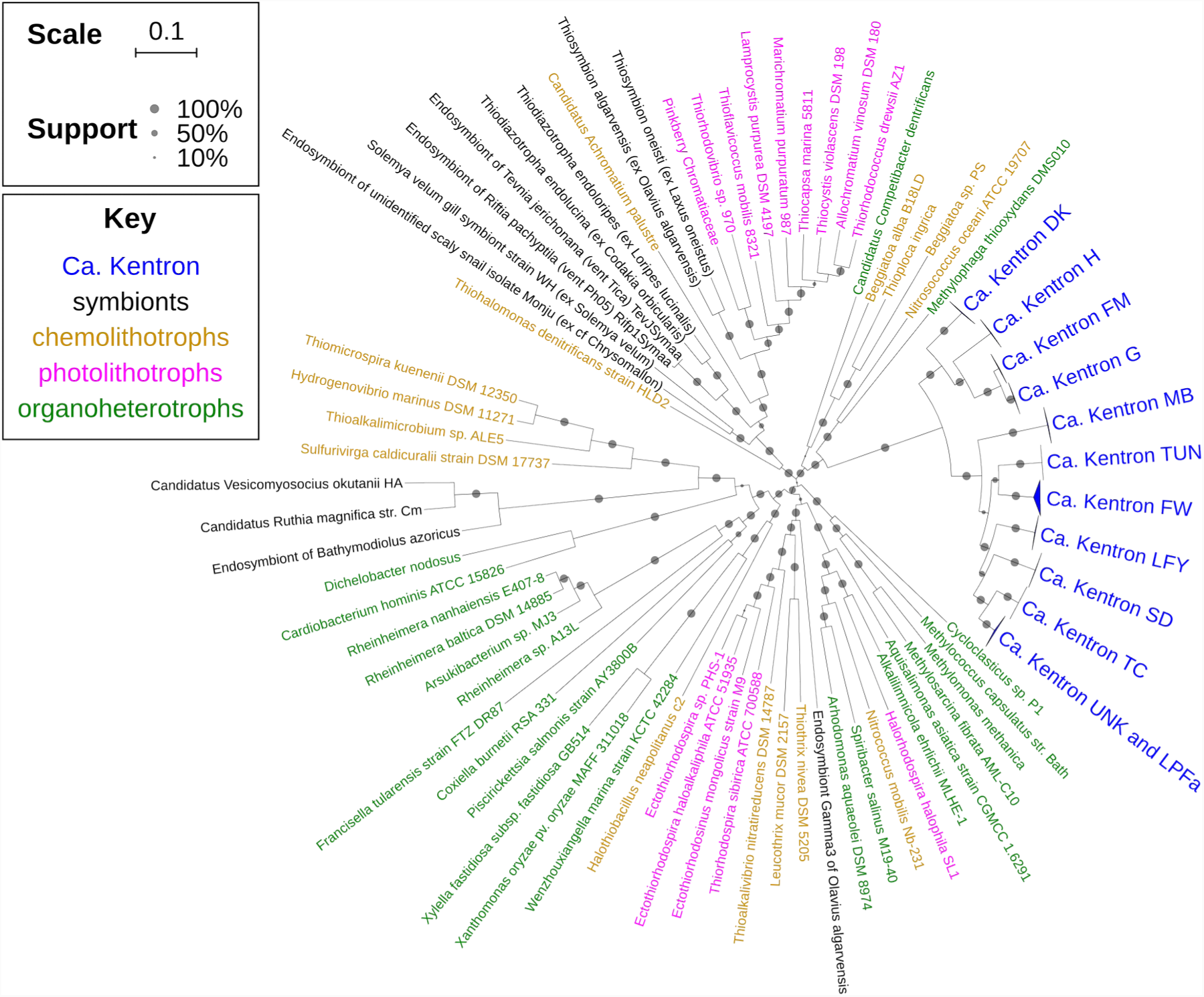
Maximum-likelihood phylogeny of Kentron and basal Gammaproteobacteria from concatenated alignment of 30 conserved protein-coding marker genes. Support values: SH-like aLRT. Branch lengths: Substitutions per site.

### Genes for key enzymes in known autotrophic pathways are absent

Unlike other investigated thiotrophic symbionts, the genes for ribulose-1,5-bisphosphate carboxylase/oxygenase (RuBisCO) and other key enzymes in known autotrophic CO_2_ fixation pathways (Supplementary Table 3) were not predicted in the binned Kentron genomes by standard annotation pipelines. A gene annotated as RuBisCO in Kentron sp. H fell within Group IV of the RuBisCO family (Supplementary Figure 3). Group IV RuBisCOs, also known as RuBisCO-like proteins (RLPs), are not known to play a role in carbon fixation but participate in a variety of other pathways such as thiosulfate metabolism (25).

To rule out the possibility that genes for these enzymes were not found because of misannotation, incomplete genome binning, or problems with genome assembly, we aligned raw, unassembled reads from *Kentrophoros* metagenome libraries to the curated SwissProt database of protein sequences. Key autotrophy proteins had coverage values (median 0.00, max 69.3 FPKM) that were always lower than the median coverage of reference proteins from the TCA and partial 3-hydroxypropionate (3HPB) pathways (Figure 2a). In 89% of cases, the coverage was at least 50-fold lower than the reference median, and if not, the majority of reads could be attributed either to other microbial genome bins in the metagenome (mostly RuBisCO or AcsB), or to to the RuBisCO-like protein in Kentron H (Supplementary Figure 4). Metatranscriptomes of two phylotypes (H and SD) were also screened with the same pipeline, and key autotrophy proteins again had coverages that were always below the median of the reference set (median 0.00, max 1.62 FPKM) (Supplementary Figure 5). We interpret this to mean that canonical autotrophy genes were indeed absent from Kentron genomes, and not merely misassembled or mispredicted.

**Figure 2.**
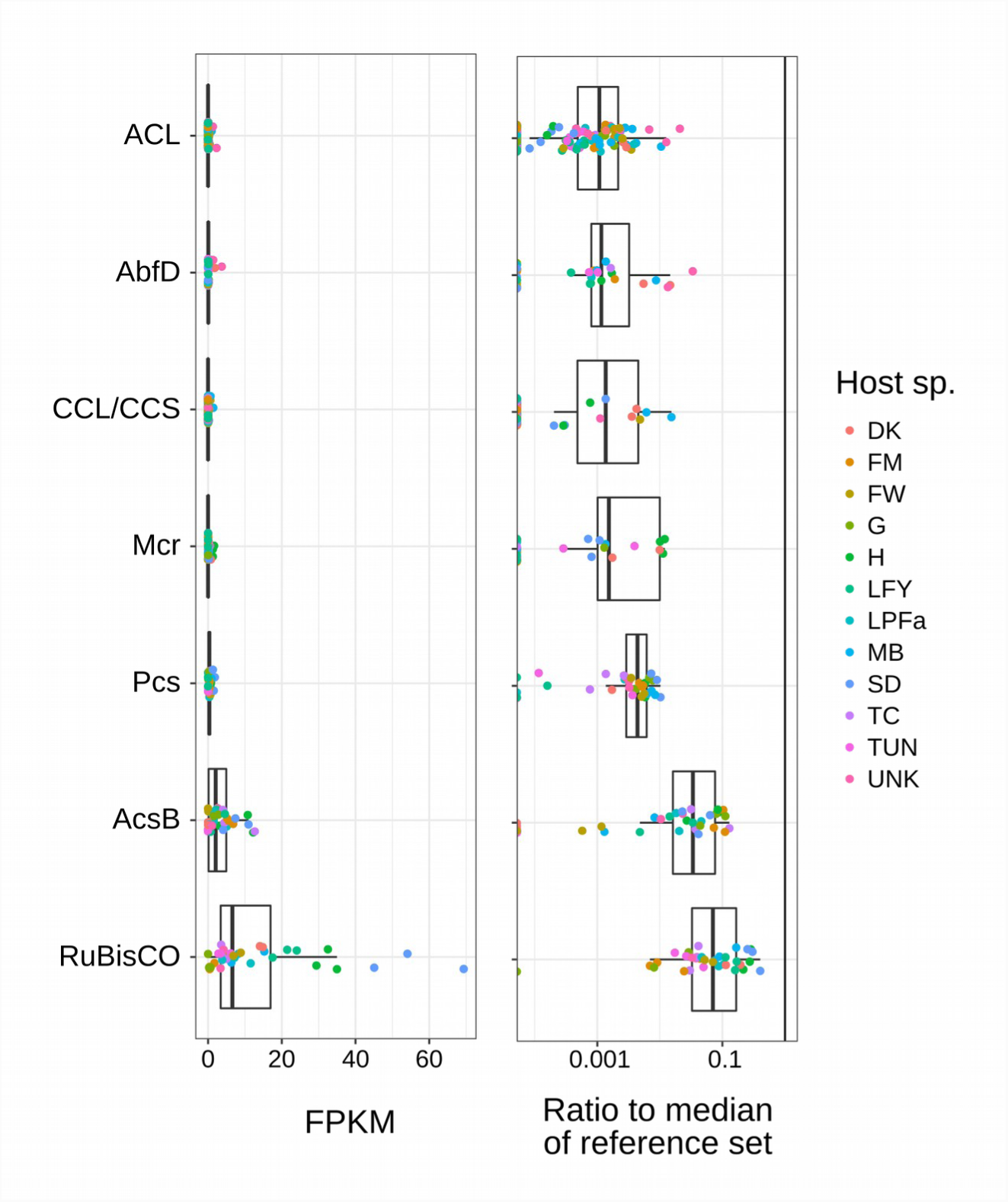
Read coverage (individual values and boxplots) in *Kentrophoros* metagenomes for key enzymes of autotrophic CO_2_-fixation pathways, expressed as FPKM values (*left*) and as a fraction of the median coverage of a reference set of proteins that are expected to be present in all Kentron species (*right*) (Supplementary Table 3). Each point represents a separate metagenome library, colored by *Kentrophoros* host morphospecies. Box midline represents median, hinges the interquartile range (IQR), whiskers are data within 1.5× IQR of hinges. *Abbreviations*: ACL, ATP citrate lyase; AbfD, 4-hydroxybutanoyl-CoA dehydratase; CCL/CCS, citryl-CoA lyase/citryl-CoA synthase; Mcr, malonyl-CoA reductase; Pcs, propionyl-CoA synthase; AcsB, CO-methylating acetyl-CoA synthase;. RuBisCO, ribulose-1,6-bisphosphate carboxylase/oxygenase.

### Evidence for lithoheterotrophic metabolism in Kentron

Kentron genome annotations suggested a lithoheterotrophic metabolism, in which energy is produced by oxidation of reduced sulfur, and carbon is assimilated in the form of organic compounds (Figure 3).

**Figure 3.**
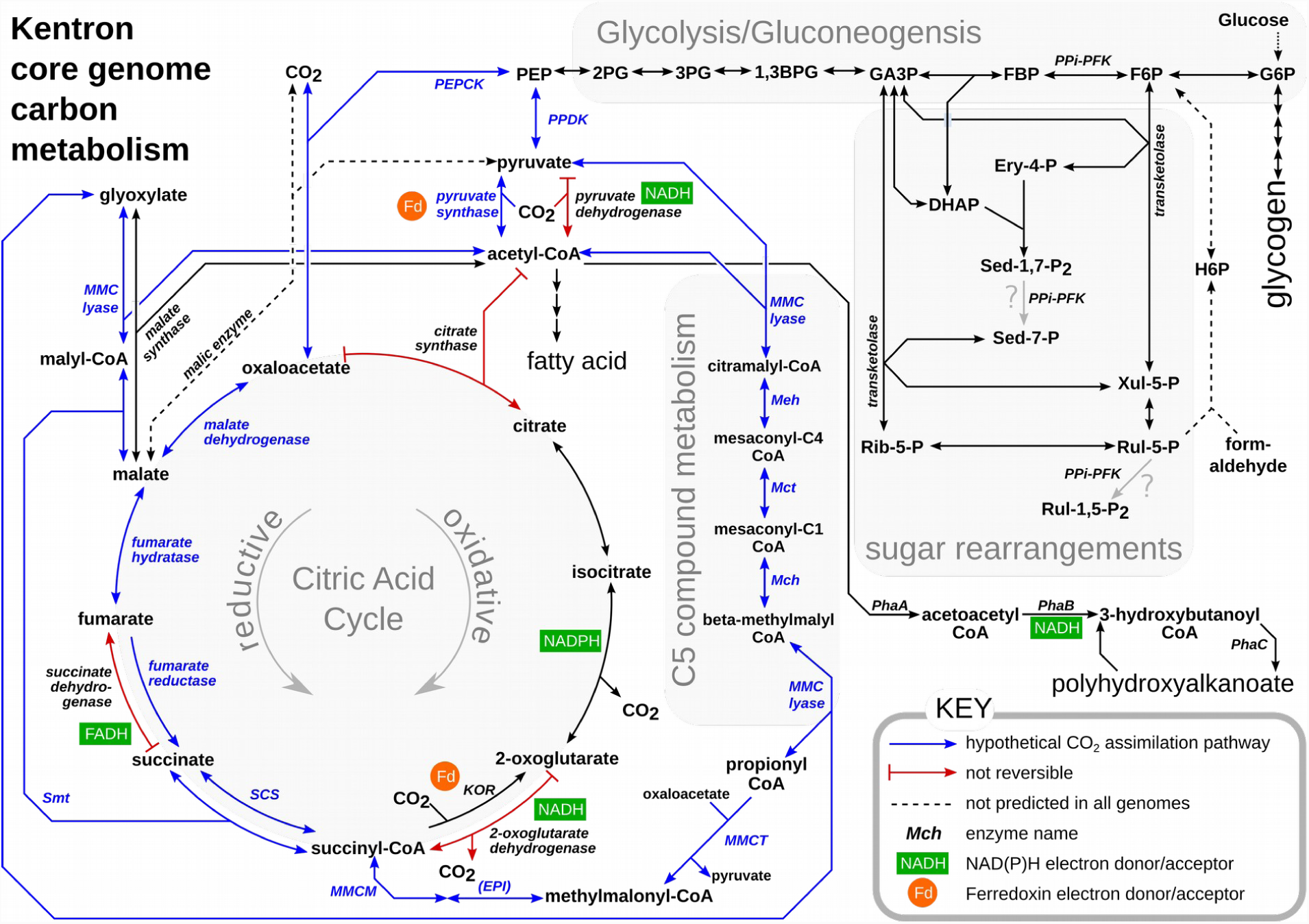
Schematic reconstruction of carbon and central metabolism of Kentron clade, focussing on pathways discussed in the text. *Compound name abbreviations*: 1,3BPG, 1,3-bisphosphoglycerate; 2PG, 2-phosphoglycerate; 3PG, 3-phosphoglycerate; DHAP, dihydroxyacetone phosphate; Ery-4-P, erythrose-4-phosphate; F6P, fructose-6-phosphate; FBP, fructose-1,6-bisphosphate; G6P, glucose-6-phosphate; GA3P, glyceraldehyde-3-phosphate; H6P, hexose-6-phosphate; PEP, phosphoenolpyruvate; Rib-5-P, ribose-5-phosphate; Rul-1,5-P2, ribulose-1,5-bisphosphate; Rul-5-P, ribulose-5-phosphate; Sed-1,7-P2, sedoheptulose-1,7-bisphosphate; Sed-7-P, sedoheptulose-7-phosphate; Xul-5-P, xylulose-5-phosphate. *Enzyme name abbreviations*: EPI, methylmalonyl-CoA epimerase; KOR, alpha-ketoglutarate oxidoreductase; Mch, mesaconyl-C1-CoA hydratase; Mct, mesaconyl-CoA C1-C4 CoA transferase; Meh, mesaconyl-C4-CoA hydratase; MMC lyase, (S)-malyl-CoA/beta-methylmalyl-CoA/(S)-citramalyl-CoA lyase; MMCM methylmalonyl-CoA mutase; MMCT, methylmalonyl-CoA carboxytransferase; PEPCK, phosphoenolpyruvate carboxykinase; PPDK, pyruvate phosphate dikinase; PPi-PFK, pyrophosphate-dependent phosphofructokinase; Smt, succinyl-CoA:(S)-malate-CoA transferase.

#### Electron donors and energetics

Kentron genomes encoded a hybrid Sox-reverse Dsr pathway for sulfur oxidation, similar to other symbiotic and free-living thiotrophs (e.g. *Allochromatium vinosum*), which would allow the use of thiosulfate, elemental sulfur, and sulfide as energy sources (26, 27). They had a complete electron transport chain for oxidative phosphorylation and an F_0_F_1_-type ATP synthase. The only terminal oxygen reductase predicted was cbb3-type cytochrome c oxidase (complex IV), which has a high oxygen affinity and is typically expressed under micro-oxic conditions (28, 29). In the two Kentron phylotypes for which expression profiles were available, this set of functions was among the most highly-expressed genes (Supplementary Figure 6).

Four Kentron phylotypes (H, SD, FW, G) encoded anaerobic-type Ni-dependent CO dehydrogenase precursors, adjacent to CO dehydrogenase Fe-S subunits (in FW and SD) or a CO dehydrogenase maturation factor (in G). In addition, H_2_ may serve as an electron donor for Kentron TC, TUN, G, and FW (one genome), which encoded genes related to the oxidative-type [Ni-Fe] hydrogenase Mvh (A and G subunits), as well as auxiliary proteins for hydrogenase maturation and Ni incorporation, although they did not all occur in a single gene cluster. Both CO and H_2_ are known to be potential electron donors for symbiotic thiotrophs, and have been measured in their habitat in Sant’ Andrea, Elba (18), where one of these *Kentrophoros* phylotypes (H) was collected.

Oxidoreductases for anaerobic respiration were not predicted, except for subunits NapA and B of periplasmic nitrate reductase (in 28 and 25 genomes respectively). However, the rest of the dissimilatory nitrate reduction to ammonia pathway was absent. Na^+^-translocating ferredoxin:NAD^+^ (Rnf) and NADH:ubiquinone (Nqr) oxidoreductases, which can couple reducing equivalents to the Na^+^ membrane potential, were also predicted.

#### Uptake transporters for organic substrates

Genes encoding uptake transporters for organic substrates were abundant in *Kentron* genomes and were also expressed in the transcriptomes (Supplementary Figure 7, Supplementary File 2). An average of 54.1 of such genes were predicted per genome (representing 18.1% of all genes with TCDB hits), of which more than half had transmembrane (TM) domains (mean 30.5 per genome). The families with the highest mean counts per genome were the ATP-binding cassette (ABC) superfamily (33.9 total, 16.4 transmembrane, counting only uptake-related subfamilies), tripartite ATP-independent periplasmic transporter (TRAP-T) family (7.2 total, 5.1 TM), and the solute:sodium symporter (SSS) family (1.6 total, 1.3 TM). Three other families – concentrative nucleoside transporter (CNT), dicarboxylate/amino acid cation symporter (DAACS), and neurotransmitter/sodium symporter (NSS) – were represented by a single gene in all Kentron genomes. Most of these families are known to target organic acids, amino acids, or small peptides. In comparison, sugar uptake transporter families were less numerous and present in only a subset of genomes (e.g. ABC subfamilies CUT 1 and CUT2), or not predicted in Kentron at all (e.g. phosphotransferase system family).

The number of organic uptake transporters in Kentron was high when compared to other symbiotic thiotrophs, which had counts ranging from 2 (0 TM) in *Ca*. Vesicomyosocius okutanii to 134 (69 TM) in the Gamma3 symbiont of *Olavius algarvensis*. However, larger genomes tend to have more transporters, and Kentron genomes were also relatively large (Figure 4a). We therefore compared the content of organic-uptake-related TCDB family members per genome between Kentron and other basal Gammaproteobacteria by non-metric multidimensional scaling. Kentron overlapped with the range of variation for both phototrophs and chemolithotrophs (both free-living and symbiotic), but were most distant from pathogenic organoheterotrophs, and from the thiotrophic symbionts of deep-sea bivalves (which have few uptake transporters) (Figure 4b).

**Figure 4.**
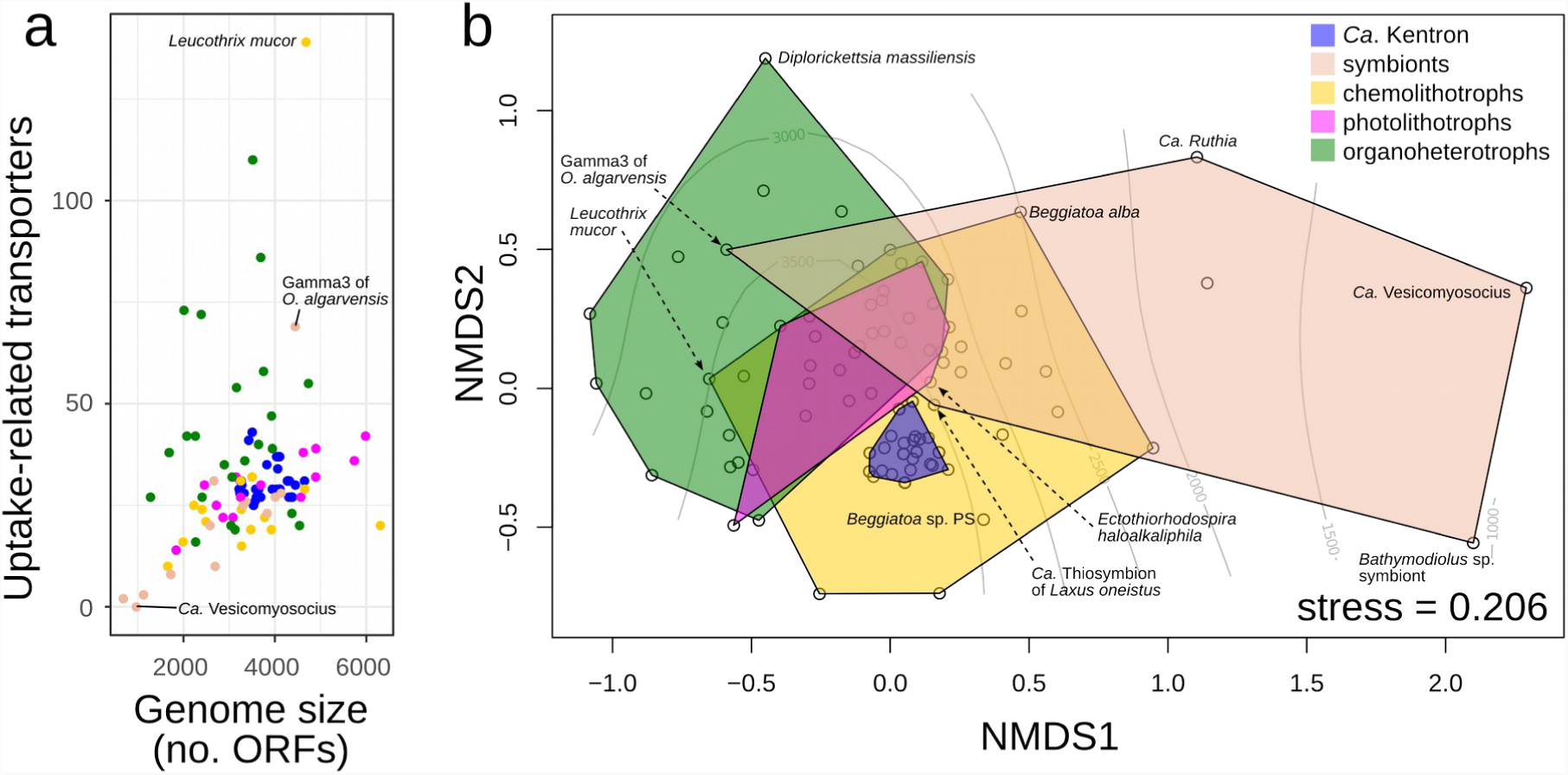
Comparison of organic substrate transporters in genomes of Kentron and other basal Gammaproteobacteria. (a) Counts of uptake-related transporters (transmembrane only) vs. genome size (expressed in no. of open reading frames). (b) 2-dimensional ordination plot (non-metric multidimensional scaling) of genomes based on counts of uptake-related TC families and subfamilies per genome. Bray-Curtis distance metric; stress = 0.206. Contour lines indicate approximate genome size. Colors in both plots share the same legend and represent type of metabolism.

#### Heterotrophic carbon metabolism

Kentron genomes encoded both glycolysis (Embden-Meyerhoff-Parnas pathway) and the oxidative tricarboxylic acid (TCA) cycle. The canonically irreversible reactions of glycolysis, pyruvate kinase and phosophofructokinase, were replaced in Kentron by pyrophosphate-dependent alternatives pyruvate phosphate dikinase and PPi-dependent phosphofructokinase (PPi-PFK) respectively. These catalyze reversible reactions that could also function in the direction of gluconeogenesis. These reversible alternatives have been found in other thiotrophic symbioses, where they have been proposed to function in a more energy-efficient version of the CBB cycle (30).

Genes for pyruvate dehydrogenase and the complete oxidative TCA cycle were present, including 2-oxoglutarate dehydrogenase, which is often missing in obligate autotrophs (31). The reductive equivalents for the key steps of the oxidative TCA cycle were also present, namely ferredoxin-dependent pyruvate synthase, ferredoxin-dependent 2-oxoglutarate synthase, and fumarate reductase. However, because neither ATP citrate lyase (ACL) nor citryl-CoA lyase/citryl-CoA synthase (CCL/CCS) were predicted, a canonical autotrophic reductive TCA cycle was not predicted.

Heterotrophic carboxylases also had relatively high expression levels. Ferredoxin-dependent pyruvate synthase was present in multiple copies per genome, of which the highest-expressed were at the 98.4 and 93.0 percentiles in Kentron H and SD respectively (Supplementary Figure 6). GDP-dependent PEP carboxykinase, which can replenish oxaloacetate anaplerotically, was also highly expressed (93.0 and 96.7 percentiles) (Supplementary Figure 6). Unlike PEP carboxylase, which was not predicted, PEP carboxykinase catalyzes a reversible reaction.

The glyoxylate shunt, which enables growth solely on acetate as the only energy and carbon source, appeared to be incomplete, as malate synthase was predicted but not isocitrate lyase. Other pathways for growth on acetate, namely the ethylmalonyl-CoA pathway and methylaspartate cycle, were not predicted either.

#### Partial 3-hydroxypropionate bi-cycle

Genes encoding most enzymes of the 3-hydroxypropionate bi-cycle (3HPB), which is the autotrophic pathway used by members of the distant bacterial phylum Chloroflexi, were predicted in Kentron. These genes had expression levels in the 64.1–83.0 and 48.2–96.4 percentile ranges for Kentron H and SD respectively (Supplementary Figure 6). The key enzymes malonyl-CoA reductase and propionyl-CoA synthase were absent, hence the bi-cycle was not closed and would not function autotrophically. However, the remainder of the pathway could function in the assimilation of organic substrates (e.g. acetate and succinate), or to connect metabolite pools (acetyl-CoA, propionyl-CoA, pyruvate, and glyoxylate) (32), as previously proposed for *Chloroflexus* (33) and the *Ca*. Thiosymbion symbionts of gutless oligochaetes (3).

These enzymes are unusual because their genes are uncommon and have a disjunct phylogenetic distribution: Chloroflexi, at least four clades in Gammaproteobacteria (Kentron, *Ca*. Thiosymbion, *Ca*. Competibacter, “Pink Berry” Chromatiaceae), and Betaproteobacteria (*Ca*. Accumulibacter). While they were previously thought to have been horizontally transferred from Chloroflexi to the other groups (33), gene phylogenies show that the Chloroflexi probably also gained the 3HPB by horizontal transfer (34), which was supported by our analysis when Kentron homologs were also included (Supplementary Figure 8).

#### Storage compounds

In addition to elemental sulfur, Kentron also have the potential to store and mobilize carbon (as polyhydroxyalkanoates (PHA) and starch/glycogen) and phosphorus (as polyphosphate). Genes related to PHA synthesis were among the most highly-expressed, namely those encoding phasin, a protein associated with the surface of PHA granules, and putative acetoacetyl-CoA reductase (*phaB*) (Supplementary Figure 6). Trehalose was detected in *Kentrophoros* sp. H but is probably produced and accumulated by the host ciliate rather than the symbionts (Supplementary Results).

### Carbon stable isotope fingerprinting (SIF) of Kentron

Measuring the natural abundance ratio of carbon stable isotopes ^13^C/^12^C, also known as the stable isotope fingerprint (SIF), is a challenge in *Kentrophoros* because of its small biomass (∼10^6^ symbionts and ∼10 µg wet weight per ciliate in the largest species). Sensitive applications of isotope ratio mass spectrometry (IRMS) for a bulk (combined host and symbionts) measurement would require at least ∼10^7^ bacterial cells (35), and compound-specific IRMS for signatures of specific pathways like ^13^C enrichment in fatty acids in the rTCA cycle (11) would require considerably more. We therefore used a newly-developed metaproteomics method that could distinguish the SIF of the symbiont from other biomass in the sample (36). The protein-based carbon SIF for Kentron sp. H from Elba and France ranged from −12.3 to −2.5 ‰ (n = 8), expressed as δ^13^C values which report deviation from the V-PDB standard (Figure 5, Supplementary Table 11). In comparison, published δ^13^C values for other shallow-water thiotrophic symbioses were < −17 ‰, and the δ^13^C of dissolved inorganic carbon (DIC) in porewater from Elba was between −2.99 and −1.32 ‰ (Figure 5, Supplementary Table 12).

**Figure 5.**
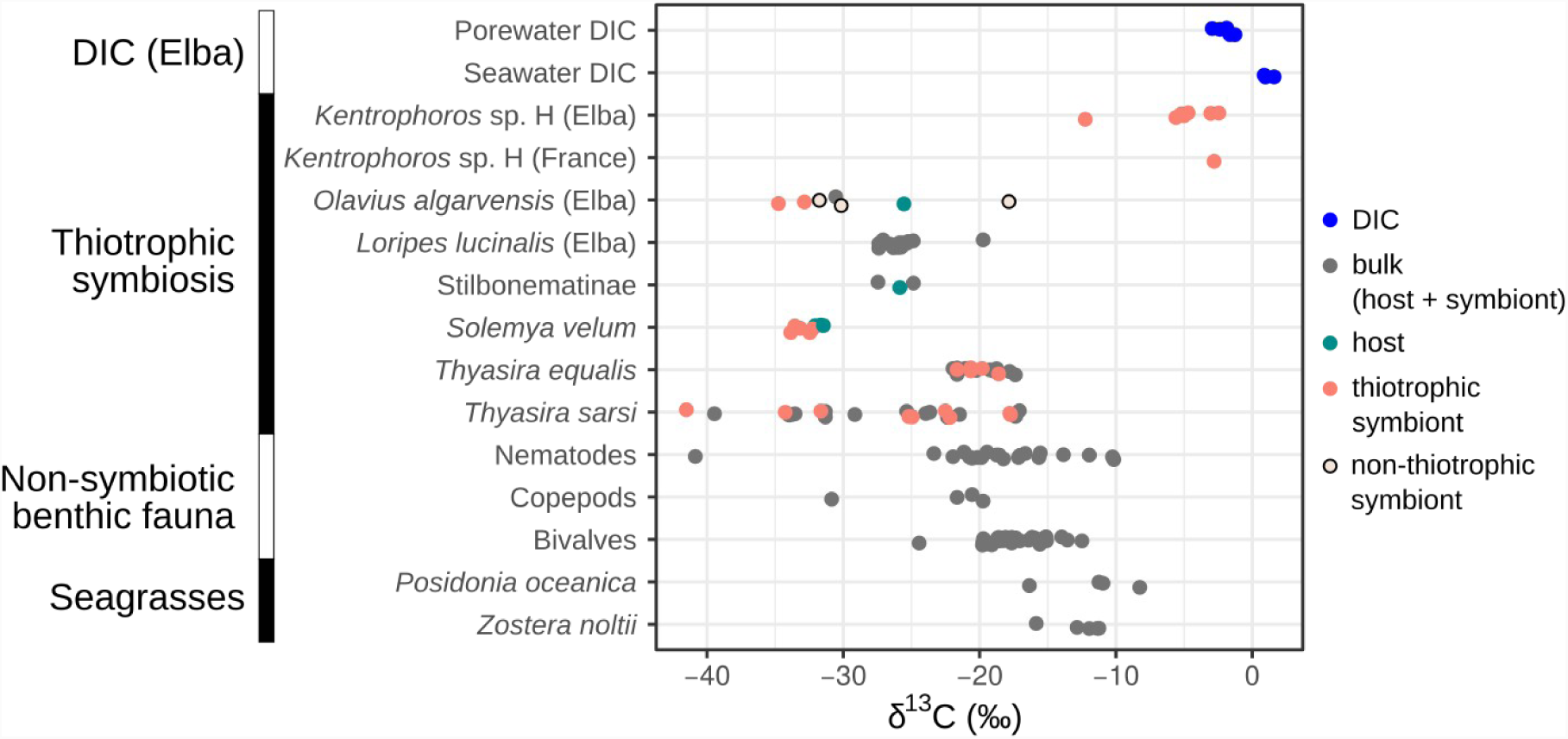
Carbon stable isotope δ^13^C composition values in *Kentrophoros* sp. H (this study) and published values for other shallow-water thiotrophic symbioses (5, 36, 43–45), non-symbiotic benthic animals (5, 58, 59), and two Mediterranean seagrass species (58–61), compared with dissolved inorganic carbon (DIC) from porewater and seawater at Elba (this study). Values for *Kentrophoros* and *Olavius algarvensis* (except the “bulk” value) are from direct protein-SIF, others are from isotope ratio mass spectrometry (IRMS). Values for symbiont-bearing tissue (e.g. gills) are also included under “symbiont”.

## Discussion

In this study, we have presented evidence that the Kentron symbionts of *Kentrophoros* ciliates are unique among thiotrophic symbionts because they do not encode canonical pathways for autotrophic carbon fixation, despite being a food source for their hosts. Their carbon stable isotope fingerprints are also substantially heavier than other thiotrophic symbioses from similar habitats. Their genomes encode heterotrophic features, including abundant uptake transporters for organic substrates, and the ability to store and mobilize organic carbon in storage polymers. We therefore propose that Kentron are chemolithoheterotrophs (37), oxidizing inorganic compounds (in this case reduced sulfur species) to provide energy for assimilating organic carbon as the main carbon source for growth.

### Role of heterotrophic CO_*2*_ *fixation*

Our results conflict with the previous interpretation of Kentron as an autotrophic symbiont, based on experiments with ^14^C-labeled bicarbonate that showed inorganic carbon fixation by Kentron at a maximum rate of 0.11 bacterial cell carbon h^-1^ (21). To rule out the possibility that only some species are autotrophs, we collected *Kentrophoros* matching the described morphology from the same site (Nivå Bay, Denmark), but their symbionts (phylotype DK) lacked canonical autotrophic pathways like all other Kentron phylotypes examined in this study.

However, the ability to fix CO_2_ alone is insufficient evidence for autotrophy, which is defined as the ability to grow with inorganic carbon as the sole or major carbon source (38), because heterotrophs can also fix CO_2_ to some extent, e.g. via anaplerotic reactions in the oxidative TCA cycle (39). Such heterotrophic fixation can account for 10% or more of total cell carbon in some bacteria (40, 41). The strictest standard of evidence for autotrophy requires cultivation to show growth in the absence of organic substrates or to measure growth rates and carbon stoichiometry, but *Kentrophoros* and its symbionts remain unculturable.

Kentron had two heterotrophic carboxylases in the central carbon metabolism, ferredoxin-dependent pyruvate synthase and PEP carboxykinase, that were both highly expressed. The former is involved in carboxylating acetyl-CoA to pyruvate, which can occur when the storage polymer PHA is mobilized. The experiments of Fenchel & Finlay (21) were performed with freshly-collected organisms that had visible cellular inclusions, and were conducted with filtered coastal seawater, which typically has more dissolved organic carbon than oceanic seawater. It is therefore likely that storage polymers and organic substrates were present in the symbiosis that were mobilized or assimilated, and that the measured CO_2_ assimilation was due to heterotrophic carboxylation.

### Could Kentron use a novel autotrophic CO_*2*_ *fixation pathway?*

Different autotrophic carbon fixation pathways each have characteristic degrees of isotope fractionation discriminating against the heavier isotope ^13^C, resulting in biomass that is relatively depleted in ^13^C (i.e. more negative δ^13^C values) (42). Kentron were more enriched in ^13^C than other shallow-water thiotrophic symbioses collected at the same locality or elsewhere, which primarily use the CBB cycle, and which have δ^13^C values in the range of −30 to −20 ‰ (Figure 5) (5, 36, 43–45). Kentron showed only a modest ^13^C depletion relative to DIC from the same site (Figure 5), which ruled out the possibility that they use a pathway with strong isotope fractionation (ε), such as the CBB cycle (ε = 10 to 22 ‰) or the reductive acetyl-CoA pathway (ε = 15 to 36 ‰) (46). Other pathways such as the reverse TCA cycle (ε = 4 to 13 ‰) or 3-hydroxypropionate bicycle (ε ≈ 0 ‰) may still fall in this range, but given that the key genes for these pathways were not detected, this possibility would require the postulation of hitherto unknown enzymes.

In two different thermophilic bacteria, the oxidative TCA cycle has recently been found to function in the autotrophic direction without using ACL or CCL/CCS, but instead by reversing the citrate synthase reaction. Such a “reversed oxidative TCA” (roTCA) cycle would not be distinguishable from the oxidative TCA by genome sequences alone (47, 48). However, both roTCA bacteria require anoxic conditions with hydrogen as the energy source for autotrophic growth. They are also facultative autotrophs, and switch to heterotrophic growth when suitable substrates like acetate are available. Citrate synthase is also highly expressed in the roTCA, whereas in Kentron the gene has only moderate expression (Supplementary Figure 6, 51.2 and 56.3 percentiles in Kentron H and SD respectively). For these reasons it is unlikely that a microaerophilic sulfur oxidizer like Kentron uses the roTCA for autotrophic growth.

Alternatively, a set of reactions that could allow autotrophic CO_2_ fixation by Kentron can be reconstructed by combining elements of the partial 3HPB and another previously proposed hypothetical pathway (49), without proposing any novel enzymes or biochemical reactions (Figure 3, Supplementary Discussion). Like the canonical 3HPB in *Chloroflexus* (33), this hypothetical pathway would allow co-assimilation of organic substrates if available, while fixing CO_2_ with ferredoxin-dependent pyruvate synthase and PEP carboxykinase, the aforementioned heterotrophic carboxylases. Although it is stoichiometrically and energetically feasible for Kentron to fix CO_2_ purely autotrophically through this hypothetical pathway, it is more likely that the involved enzymes function lithoheterotrophically or mixotrophically, enabling them to exploit different carbon sources at the same time (Supplementary Discussion).

### The autotrophy-heterotrophy spectrum in thiotrophic symbiosis

Thiotrophic symbioses are most commonly found in nutrient-limited environments, and their symbionts are assumed to provide the hosts with nutrition through the autotrophic fixation of CO_2_. Indeed, the symbionts of deep-sea bivalves *Bathymodiolus* and *Calyptogena* show characteristic features of obligate autotrophy in their genomes, namely an incomplete TCA cycle and the lack of organic uptake transporters (Table 1) (19, 50–52). This appears to be the exception, however, as other symbiont clades possess heterotrophic features to varying degrees (Table 1). Some features, e.g. glycolysis, are involved in the mobilization of storage compounds, but abundant presence and expression of organic uptake transporters, as we observed in Kentron in this study, are a clearer marker of heterotrophic assimilation (53). Mixotrophic potential in other symbionts has been variously suggested to be a strategy to cope with carbon limitation by recycling host waste, as a nutritional supplement to autotrophy, or to be retained for a hypothetical free-living stage of the symbiont life cycle (3, 7, 30). Thus, there is a spectrum among thiotrophic symbionts between obligate autotrophs and the possibly heterotrophic Kentron, with various degrees of mixotrophy in between.

**Table 1.** Comparison of metabolic features predicted in thiotrophic symbiont genomes. *Key*: +, present; (+), not in all genomes. *Abbreviations*: Bathy, *Bathymodiolus;* CBB, Calvin-Benson-Bassham cycle; Cyt c, Cytochrome c; Frd, fumarate reductase; Kor, 2-oxoglutarate:ferredoxin oxidoreductase; PEP, phosphoenolpyruvate; PPi, pyrophosphate; rTCA, reverse TCA cycle.

| Feature | Bathy<br>symbiont | Ruthia | Endoriftia | Thio-<br>diazotropha | Solemya<br>symbiont | Thio-<br>symbion | Kentron |
| --- | --- | --- | --- | --- | --- | --- | --- |
| Host habitat | Hydrothermal vents and seeps |  |  | Shallow water sediment interstitial |  |  |  |
| Autotrophy |  |  |  |  |  |  |  |
| CBB cycle (RuBisCO, phosphoribulokinase) | + | + | + | + | + | + |  |
| PPi-Phosphofructokinase | + | + | + | + | + | + | + |
| rTCA (Citrate cleavage) |  |  | + |  |  |  |  |
| Diazotrophy |  |  |  |  |  |  |  |
| Nitrogenase |  |  |  | + |  | (+) |  |
| Tricarboxylic acid (TCA) cycle |  |  |  |  |  |  |  |
| Oxidative TCA |  |  | + | + | + | + | + |
| Kor & Frd (reductive TCA) |  |  | + | + |  | + | + |
| Glyoxylate shunt |  |  |  | + | + | + |  |
| Central metabolism |  |  |  |  |  |  |  |
| Pyruvate phosphate dikinase |  |  | + | + | + | + | + |
| PEP synthase |  |  | + | + | + | + |  |
| Pyruvate synthase |  |  | + | + | + | + | + |
| Pyruvate carboxylase |  |  |  | + |  | + |  |
| PEP carboxylase |  |  |  | + | + |  |  |
| PEP carboxykinase (GTP) |  |  |  | + |  | + | + |
| Malic enzyme | + | + | + | + | + | + | (+) |
| C5 reactions of 3-hydroxypropionate bi-cycle (p3HPB) |  |  |  |  |  |  |  |
| p3HPB |  |  |  |  |  | + | + |
| Energy |  |  |  |  |  |  |  |
| Rnf transporter | + | + | + | + | + | + | + |
| V-type ATPase |  |  | + | + | + |  |  |
| Cyt c oxidase cbb3 type | + | + | + | + | + | + | + |
| Cyt c oxidase aa3 type | + | + |  | + | + | + |  |
| Storage compounds |  |  |  |  |  |  |  |
| Glycogen |  |  | + | + | + | + | + |
| Polyhydroxyalkanoates |  |  |  | + | + | + | + |
| Polyphosphate synthesis |  |  |  | + | + | + | + |

Symbiotic thiotrophs that lack the canonical CBB and rTCA pathways, as Kentron does, have not been previously described. Among free-living thiotrophic bacteria, lithoheterotrophy appears to be more common among those that have the Sox pathway (i.e. thiosulfate oxidizers) than those with the rDsr/Sox pathway (i.e. thiotrophs that can store and oxidize elemental sulfur) (Supplementary Discussion). Of the latter, we are aware of two isolates – *Ruegeria marina* CGMCC 1.9108 and *Thiothrix flexilis* DSM 14609 – whose genomes lack CBB and rTCA. Moreover, some free-living thiotrophs that possess the CBB cycle may nonetheless grow only when supplied with organic substrates, e.g. freshwater *Beggiatoa* (54). Functional heterotrophy may therefore be underestimated as it is not necessarily apparent from genomic predictions.

Host biology constrains the feasibility of autotrophy for a thiotrophic symbiont. To meet nutritional requirements by chemoautotrophy alone, the host must provide high O_2_ flux to its symbionts, beyond what it requires itself (55). This is metabolically demanding, and it is telling that the bathymodioline and vesicomyid bivalves, whose symbionts have the most autotrophic features, are relatively large animals with intracellular symbionts that are located in their gill tissues, which can better maintain ventilation and homeostasis than smaller hosts that have extracellular symbionts. Specialization for high autotrophic production rates is also seen in the pre-concentration of CO_2_ by the bivalve *Bathymodiolus azoricus* for its symbionts, and in its thiotrophic symbiont’s metabolic dependence on the animal to replenish TCA cycle intermediates (56).

Meiofaunal hosts like *Kentrophoros* and stilbonematine nematodes, in contrast, are much smaller, cannot span substrate gradients, and must be able to tolerate fluctuating anoxia. Given that shallow-water coastal environments also receive more organic input, for example from land or from seagrass beds, than deep-sea hydrothermal environments, it is not surprising that the shallow-water meiofaunal symbioses have more heterotrophic features than the deep-sea ones (Table 1).

### Ecophysiological model of the Kentrophoros *symbiosis*

Based on our results and previous descriptions of morphology and behavior in *Kentrophoros* and other thiotrophic symbioses, we propose the following model for the ecophysiology of this symbiosis:

*Kentrophoros* fuels its growth by the phagocytosis and digestion of its symbionts, which was previously observed by electron microscopy (22). There has to be a net input of energy and organic carbon from environmental sources for the overall growth of the host-symbiont system, and heterotrophic carboxylation may also be a substantial carbon source. To give its symbionts access to these substrates, *Kentrophoros* likely shuttles between oxic and anoxic zones in marine sediment, like other motile, sediment-dwelling hosts with thiotrophic symbionts (43). In anoxic sediment, both the predicted energy and carbon sources, namely sulfide and organic acids, are produced by microbial activity (57). Many organic acids, such as acetate and succinate, are more oxidized than average biomass (Supplementary Table 4), hence Kentron needs reducing equivalents to assimilate them, which could come from sulfide. As the complete oxidation of sulfide to sulfate requires oxygen, the partly-oxidized sulfur can be stored by the symbionts as elemental sulfur when under anoxic conditions, until the symbionts are again exposed to oxygen. The synthesis of PHA from small organic acids like acetate can also function as both an additional electron sink for sulfide oxidation and as a carbon store. Hydrolysis of polyphosphate and mobilization of glycogen are also potential sources of energy in the absence of oxygen.

Under oxic conditions, elemental sulfur inclusions in Kentron can be further oxidized to sulfate to yield energy, and PHA can be mobilized for biosynthesis. Glycogen and polyphosphate reserves can also be regenerated. The various storage inclusions in Kentron, namely elemental sulfur, PHA, glycogen, and polyphosphate, hence, represent pools of energy, reducing equivalents, and carbon that function as metabolic buffers for the symbiont living in a fluctuating environment.

The symbionts may also bring a syntrophic benefit to their hosts under anoxic conditions, when the ciliates can only yield energy by fermentation. By assimilating fermentation waste products and keeping their concentrations low in their host, the symbionts can improve the energy yields for their hosts and allow them to better tolerate periods of anoxia. This could also be a form of resource recycling under carbon-limited conditions, which has been proposed for other thiotrophic symbionts with the potential to assimilate organic acids (3, 7). Kentron is relatively enriched in ^13^C compared to non-symbiotic shallow-water benthic fauna, such as nematodes and bivalves (δ^13^C ∼ −20 to −10 ‰) (5, 58, 59), and to the seagrasses (δ^13^C ∼ −15 to −10 ‰) (58–61) that are the main primary producers in the habitat of *Kentrophoros* (Figure 5). The higher values in Kentron could be partly caused by preferring specific substrates with higher ^13^C content, such as acetate, which has a wide range of δ^13^C (−2.8 to −20.7 ‰) in marine porewaters depending on the dominant microbial processes at the site (62). Given how close the δ^13^C of Kentron is to DIC, it is possible that heterotrophic CO_2_ fixation contributes to this ^13^C signature, but the isotope fractionation values of the heterotrophic carboxylases have not been characterized, to our knowledge. Repeated internal recycling of host waste products, as we postulate, could also cause accumulation of ^13^C in the host-symbiont system.

Our metabolic model has parallels to free-living thiotrophs (63, 64) and to heterotrophic bacteria involved in enhanced biological phosphorus removal (EBPR) from wastewater (65). What they have in common is their use of storage inclusions as metabolic buffers for fluctuating oxygen and nutrient conditions. For example, lithomixotrophic giant sulfur bacteria like *Thiomargarita* and *Thioploca* survive anoxia by using nitrate stored in vacuoles as an alternative electron acceptor to partially oxidize sulfide to elemental sulfur. They also use polyphosphate for energy and can store assimilated carbon as glycogen or PHA (64, 66).

### Conclusion

We have shown that a diverse and widespread clade of symbiotic sulfur bacteria lacks genes encoding canonical enzymes for autotrophic CO_2_ fixation, despite being a food source for their hosts. This is unlike all other thiotrophic symbionts sequenced to date, which possess the CBB or rTCA cycles for autotrophy. We propose a lithoheterotrophic model for the *Kentrophoros* nutritional symbiosis, which challenges the chemoautotrophic paradigm usually applied to thiotrophic symbiosis. Uptake of organic substrates from the environment, heterotrophic carboxylation, and recycling of host waste may play a bigger part in thiotrophic symbioses than previously thought. Our results suggest that nutritional symbioses can also be supported by chemolithoheterotrophy, and that thiotrophic symbioses fall on a spectrum between autotrophy and heterotrophy.

## Materials and Methods

### Sample collection

Specimens of *Kentrophoros* were collected in 2013 and 2014 from Elba, Italy (Mediterranean Sea), in 2015 from Twin Cayes, Belize (Caribbean Sea), and in 2016 from Nivå Bay, Denmark (Øresund Strait between Baltic and North Sea), as previously described (20). Sampling localities and dates, as well as the number of specimens and phylotypes that were sequenced are given in Supplementary Table 1.

### DNA/RNA extraction and sequencing

Samples for DNA and RNA extraction, comprising single ciliate cells and their symbionts, were fixed in RNAlater (Ambion) and stored at 4 °C. Before DNA extraction, samples were centrifuged (8000 g, 5 min) and excess RNAlater was removed by pipetting. DNA was extracted with the DNeasy Blood and Tissue kit (Qiagen) following manufacturer’s instructions, and eluted in 50 µL of buffer AE. DNA concentration was measured fluorometrically with the Qubit DNA High-Sensitivity kit (Life Technologies). Each DNA sample was screened by PCR with eukaryotic 18S rRNA primers EukA/EukB (67) followed by capillary sequencing to identify the *Kentrophoros* phylotype, as previously described (20). Libraries for metagenomic sequencing were prepared with the Ovation Ultralow Library System V2 kit (NuGEN) following manufacturer’s protocol. Libraries were sequenced as either 100 or 150 bp paired-end reads on the Illumina HiSeq 2500 platform.

RNA was extracted with the RNeasy Plus Micro Kit (Qiagen) following manufacturer’s protocol, and eluted in 15 µL RNase-free water. cDNA was synthesized with the Ovation RNASeq System v2 (NuGEN) following manufacturer’s protocol, sheared to 350 bp target size with Covaris microTUBE system, cleaned up with Zymo Genomic DNA Clean & Concentrator Kit. Sequencing library was prepared from cDNA with NEBNext Ultra DNA library preparation kit for Illumina, and sequenced on the Illumina HiSeq 2500 platform as 100 bp single-end reads.

Library preparation and sequencing were performed at the Max Planck Genome Centre Cologne, Germany (http://mpgc.mpipz.mpg.de/home/).

### Assembly, binning, and annotation of symbiont genomes

Reads were trimmed from both ends to remove fragments matching Truseq adapters, and to remove bases with Phred quality score < 2, using either Nesoni v0.111 (https://github.com/Victorian-Bioinformatics-Consortium/nesoni) or BBmap v34+ (https://sourceforge.net/projects/bbmap/). Trimmed reads were error-corrected with BayesHammer (68). Error-corrected reads were assembled with IDBA-UD v1.1.1 (69) or SPAdes v3.5.0+ (68) to produce the initial assembly. The reference coverage of each contig was obtained by mapping the error-corrected read set against the assembly with BBmap (“fast” mode). Conserved marker genes in the assembly were identified and taxonomically classified with Amphora2 (70) or Phyla-Amphora (71). 16S rRNA genes were identified with Barrnap v0.5 (https://github.com/tseemann/barrnap) and classified by searching against the Silva SSU-Ref NR 119 database (72) with Usearch v8.1.1831 (73). Differential coverage information (74) was obtained by mapping reads from other samples of the same host morphospecies onto the assembly with BBmap. Contigs belonging to the primary *Kentrophoros* symbiont (the “primary symbiont bin”) were heuristically identified by a combination of differential coverage, assembly graph connectivity, GC%, affiliation of conserved marker genes, and affiliation of 16S rRNA sequence using gbtools v2.5.2 (75). Reads mapping to the primary symbiont bin were reassembled with SPAdes. Binning and reassembly of the primary symbiont genome was iteratively repeated for each metagenome sample until the primary symbiont bin appeared to contain only a single genome, based on the number and taxonomic affiliation of conserved marker genes and 16S rRNA. For final genome bins, summary statistics were computed with Quast v4.4 (76), and completeness and contamination were estimated with CheckM v1.0.11 (77) using the Gammaproteobacteria taxonomy workflow. Average amino-acid identity (AAI) and average nucleotide identity (ANI) values between genomes were calculated with CompareM v0.0.21 (https://github.com/ dparks1134/CompareM) and jSpecies v1.2.1 respectively (78).

Genome bins were annotated with the IMG/M pipeline for downstream analyses (79). Metabolic pathways were predicted from the annotated proteins with the PathoLogic module (80) of Pathway Tools v20.5 (81), followed by manual curation. Metabolic modules from KEGG (82) were also predicted with the KEGG Mapper tool (http://www.kegg.jp/kegg/mapper.html, accessed Jan 2017) from KEGG Orthology terms in the IMG annotation.

### Core-and pan-genome analysis

Ortholog clusters of Kentron protein sequences were predicted by first performing a reciprocal Blastp (version 2.2.29+) search (83) of all translated open reading frames (ORFs) annotated by the IMG pipeline (E-value cutoff 10^-5^), and then identifying clusters in the search results with the Markov cluster algorithm (84) using FastOrtho (inflation value 1.5), which is a reimplementation of OrthoMCL (85) by the PATRIC project (86). Accumulation curves and uncertainty estimates for the core and pan genome size were generated by random resampling (n = 200) of genome memberships for the predicted orthologs.

### Transcriptome analysis

Metatranscriptome reads for *Kentrophoros* sp. H and SD were mapped on to symbiont genome assemblies from the respective species (IMG genome IDs 2609459750 and 2615840505) using BBmap (minimum identity 0.97). Read counts per genomic feature were calculated with featureCounts v1.5.2 (87), and transformed into FPKM values (fragments per kbp reference per million reads mapped).

### Verifying absence of key genes for autotrophic pathways

Key enzymes that are diagnostic for known autotrophic pathways were identified from the literature (42, 88–90) (Supplementary Table 3). These were absent from Kentron genome annotations, with the exception of a RuBisCO-like protein (RLP) in Kentron sp. H (see below). To verify that the absence of autotrophy-related sequences was not caused by incomplete genome bins, misprediction of open reading frames, or misassembly of the reads, we aligned raw reads from host-symbiont metagenomes and metatranscriptomes against the UniProt SwissProt database (release 2017_01) (91) using diamond blastx (v0.8.34.96, “sensitive” mode) (92). Sequences for certain key enzymes were absent from SwissProt, so representative sequences from UniProtKB were manually added to the database (Supplementary Table 3, Supplementary File 4). Reads with hits to target enzymes (identified by EC number or from the list of additional sequences) were counted, extracted, and mapped against the initial metagenomic assembly for the corresponding library. Raw counts of reads were transformed to FPKM values using three times the mean amino acid length of the target proteins as the reference length. As a comparison, FPKM values were also calculated for a reference set of enzymes of the TCA cycle and partial 3HPB pathway (Supplementary Table 3), which were annotated in Kentron genomes and thus expected to have much higher coverage than the putatively absent genes.

### Identification of transporter genes for substrate uptake

Families and subfamilies of transporter proteins from the Transporter Classification Database (TCDB, accessed 2 Feb 2017) (93) that were described as energy-dependent uptake transporters for organic substrates were shortlisted (Supplementary Table 5). Translated ORFs for Kentron and selected genomes of other symbiotic and free-living basal Gammaproteobacteria (Supplementary Table 6) were aligned with Blastp (83) (best-scoring hit with E-value < 10^-5^, >30% amino acid sequence identity, and >70% coverage of reference sequence, parameters from (53)) against TCDB. As TCDB also includes non-membrane proteins that are involved in transport (e.g. ATPase subunit of ABC transporters), we also counted how many hits contained transmembrane domains, predicted with tmhmm v2.0c (94). To compare the transporter content between genomes, the tabulated counts of organic substrate uptake TC family hits per genome were analyzed by non-metric multidimensional scaling (NMDS) with the metaMDS function in the R package vegan v2.5.1 (https://CRAN.R-project.org/package=vegan) (Bray-Curtis distance, 2 dimensions, 2000 runs).

### Phylogenetic analyses

Maximum-likelihood phylogenetic trees were inferred from the following alignments with Fasttree v2.1.7 (95), using the JTT model with CAT approximation (20 rate categories) and SH-like support values.

#### Kentron and related Gammaproteobacteria

Conserved marker genes from Kentron and selected basal Gammaproteobacteria (Supplementary Table 6) were extracted by the Amphora2 pipeline. Amino acid sequences of 30 markers were aligned with Muscle v3.8.31 (96) and concatenated.

#### RuBisCO-like protein from Kentron sp. H

RuBisCO superfamily protein accessions and their classification were obtained from (25). These were aligned with RuBisCO-like protein sequences from Kentron sp. H and RuBisCO from selected sulfur-oxidizing symbiotic Gammaproteobacteria, using Muscle.

#### Proteins of partial 3-hydroxypropionate bi-cycle

Homologs to proteins of the 3-hydroxypropionate bi-cycle in *Chloroflexus aurantiacus* were obtained from the UniRef50 clusters containing the *C. aurantiacus* sequences in the UniProt database. These were aligned with the Kentron homologs with Muscle.

### Protein extraction and peptide preparation

Samples of *Kentrophoros* sp. H for proteomics were collected by decantation from sediment adjacent to seagrass meadows at Sant’ Andrea, Isola d’Elba, Italy on 3 June 2014, and from Pampelonne Beach, Provence-Alpes-Côte d’Azur, France in July 2018. Ciliates were individually fixed in RNAlater, and subsequently stored at 4 °C and then at −80 °C. One individual *Kentrophoros* sp. H specimen and nine pooled samples of four or five individuals each (Supplementary Table 11) were used to prepare tryptic digests following the filter-aided sample preparation (FASP) protocol (97) with minor modifications (98). Samples were lysed in 30 µl of SDT-lysis buffer (4% (w/v) SDS, 100 mM Tris-HCl pH 7.6, 0.1 M DTT) by heating to 95 °C for 10 min. To avoid sample losses we did not clear the lysate by centrifugation after lysis. Instead, we loaded the whole lysate on to the 10 kDa filter units used for the FASP procedure. The Qubit Protein Assay Kit (Thermo Fisher Scientific, Life Technologies) was used to determine peptide concentrations, following the manufacturer’s instructions. Peptide concentrations were below the detection limit in all samples.

### 1D-LC-MS/MS

All peptide samples were analyzed by 1D-LC-MS/MS as previously described (99), with the modification that a 75 cm analytical column was used. Briefly, an UltiMate 3000 RSLCnano Liquid Chromatograph (Thermo Fisher Scientific) was used to load peptides with loading solvent A (2% acetonitrile, 0.05% trifluoroacetic acid) onto a 5 mm, 300 µm ID C18 Acclaim PepMap100 pre-column (Thermo Fisher Scientific). Since peptide concentrations were very low, complete peptide samples (80 µL) were loaded onto the pre-column. Peptides were eluted from the pre-column onto a 75 cm × 75 µm analytical EASY-Spray column packed with PepMap RSLC C18, 2 µm material (Thermo Fisher Scientific) heated to 60° C. Separation of peptides on the analytical column was achieved at a flow rate of 225 nL min^-1^ using a 460 min gradient going from 98% buffer A (0.1% formic acid) to 31% buffer B (0.08% formic acid, 80% acetonitrile) in 363 min, then to 50% B in 70 min, to 99% B in 1 min and ending with 26 min 99% B. Eluting peptides were analyzed in a Q Exactive Plus hybrid quadrupole-Orbitrap mass spectrometer (Thermo Fisher Scientific). Carryover was reduced by running two wash runs (injection of 20 µL acetonitrile) between samples. Data acquisition in the Q Exactive Plus was done as previously described (5).

### Protein identification and quantification

A database containing protein sequences predicted from the *Ca*. Kentron genomes described above and predicted protein sequences from a preliminary host transcriptome was used for protein identification. The *Ca*. Kentron protein sequences were clustered at 98% identity with CD-HIT v4.7 (100), and only the representative sequences were used for the protein identification database. The cRAP protein sequence database (http://www.thegpm.org/crap/), which contains sequences of common lab contaminants, was appended to the database. The final database contained 5,715 protein sequences. For protein identification, MS/MS spectra were searched against this database using the Sequest HT node in Proteome Discoverer version 2.2 (Thermo Fisher Scientific) as previously described (5).

### Direct Protein-SIF

Stable carbon isotope fingerprints (SIFs = δ^13^C values) for *Ca*. Kentron symbiosis were determined using the proteomic data (36). Briefly, human hair with a known δ^13^C value was used as reference material to correct for instrument fractionation. A tryptic digest of the reference material was prepared as described above and analyzed with the same 1D-LC-MS/ MS method as the samples. The peptide-spectrum match (PSM) files generated by Proteome Discoverer were exported in tab-delimited text format. The 1D-LC-MS/MS raw files were converted to mzML format using the MSConvertGUI available in the ProteoWizard tool suite (101). Only the MS^1^ spectra were retained in the mzML files and the spectra were converted to centroided data by vendor algorithm peak picking. The PSM and mzML files were used as input for the Calis-p software (https://sourceforge.net/projects/calis-p/) to extract peptide isotope distributions and to compute the direct Protein-SIF δ^13^C value for *Ca*. Kentron and the human hair reference material (36). The direct Protein-SIF δ^13^C values were corrected for instrument fragmentation by applying the offset determined by comparing the direct Protein-SIF δ^13^C value of the reference material with its known δ^13^C value. We obtained between 50 and 499 peptides with sufficient intensity for direct Protein-SIF from seven of the nine pooled samples (Supplementary Table 11). These samples were thus well above the necessary number of peptides needed to obtain an accurate estimate. Due to the low biomass of the individual *Kentrophoros* specimen (∼ 10 µg) only 14 peptides with sufficient intensity for direct Protein-SIF were obtained for this sample. This lower number of peptides for the individual specimen can potentially lead to a lower accuracy of the respective SIF value, however, since the value fell in the same range as for the pooled samples we assume that the estimate is sufficiently accurate.

### Dissolved inorganic carbon δ^*13*^*C*

Seawater and porewater samples were collected from the vicinity of seagrass meadows at Sant’ Andrea, Elba, Italy in July 2017 to determine the δ^13^C of dissolved inorganic carbon (DIC). Seawater was sampled at the surface from a boat, whereas porewater was sampled at 15 cm sediment depth with a steel lance. Samples were drawn into 20 mL plastic syringes; 6 mL of each was fixed with 100 µL of 300 mM ZnCl_2_, and stored at 4 °C until processing. δ^13^C was measured with a Finnigan MAT 252 gas isotope ratio mass spectrometer with Gasbench II (Thermo Scientific), using Solnhofen limestone as a standard and 8 technical replicates per sample.

### Data availability

Annotated genomes are available on the Joint Genome Institute GOLD database (https://gold.jgi.doe.gov/) under study Gs0114545. Metagenomic and metatranscriptomic sequence libraries are deposited in the European Nucleotide Archive under study accessions PRJEB25374 and PRJEB25540 respectively. The mass spectrometry metaproteomics data, direct Protein-SIF relevant files, and protein sequence database have been deposited to the ProteomeXchange Consortium via the PRIDE partner repository (https://www.ebi.ac.uk/pride/archive/) with the dataset identifier PXD011616. [Reviewer access: username –, password – eLmqKA0b.] Supplementary files are available via Zenodo at doi:10.5281/zenodo.2555833.

### Code availability

Scripts used to screen for autotrophy-related genes in metagenome libraries, to classify transporter families, and to calculate phylogenetic trees are available at https://github.com/kbseah/mapfunc, https://github.com/kbseah/tcdbparse_sqlite, and https://github.com/kbseah/phylogenomics-tools respectively.

## Acknowledgements

We thank, for hosting and coordinating field work, staff of the HYDRA Institute on Elba, especially Miriam Weber, Hannah Kuhfuß, and Matthias Schneider; the Smithsonian CCRE program and staff at the Carrie Bow Caye field station; Lasse Riemann and the Marine Biological Section of the University of Copenhagen. Mario Schimak, Oliver Jäckle, Juliane Wippler, Judith Zimmermann, Miriam Sadowski, Silke Wetzel, Nikolaus Leisch, and Anne-Christin Kreutzmann assisted in sample collection. Library preparation and sequencing was performed at the Max Planck Genome Centre Cologne. We thank Marc Strous for access to proteomics equipment. The proteomics and direct Protein-SIF work was supported by the Campus Alberta Innovation Chair Program, the Canadian Foundation for Innovation, and a discovery grant from the Natural Sciences and Engineering Research Council (NSERC) of Canada (above to Marc Strous), and the NC State Chancellor’s Faculty Excellence Program Cluster on Microbiomes and Complex Microbial Communities (MK). We thank Dolma Michellod and Alexander Gruhl for collecting DIC samples, and Henning Kuhnert and the MARUM Stable Isotope Laboratory team for DIC IRMS measurements. We also thank Roland Dieterich for computational support, Caitlin Petro, Erik Puskas, Tora Gulstad, and Frantisek Fojt for mass spectrometry support, and Lizbeth Sayavedra, Oliver Müller-Caja, Monika Bright, Jörg Ott, Verena Carvalho, Cameron Callbeck, Marc Mußmann, and members of the Symbiosis Department for useful discussions and comments. Financial support was provided by the Max Planck Society, the Humboldt Foundation to CPA, Marie Curie Fellowship to HGV, the Gordon and Betty Moore Foundation Marine Microbial Initiative Investigator Award to ND (Grant GBMF3811), SFB 987 “Microbial Diversity in Environmental Signal Response” to LSvB and TJE, and FET-Open Grant 686330 (‘FutureAgriculture’) to JZ. Contribution XXX from the Caribbean Coral Reef Ecosystems (CCRE) Program, Smithsonian Institution.

## Author contributions

BS, ND, HGV designed study. BS, HGV performed field work. BH prepared sequencing libraries with BS and coordinated sequencing. ML performed metabolomics mass spectrometry analyses. AK prepared samples and generated data for proteomics. MK and AK processed and analyzed proteomics data. MK performed protein-SIF analysis. BS, CPA, JZ, LSvB, TJE, ML, HGV analyzed genomics and transcriptomics data. BS wrote manuscript draft. All authors participated in revising manuscript.

## References

1. Dubilier N, Bergin C, Lott C (2008) Symbiotic diversity in marine animals: the art of harnessing chemosynthesis. Nat Rev Microbiol 6(10):725–740.

2. Stewart FJ, Newton ILG, Cavanaugh CM (2005) Chemosynthetic endosymbioses: adaptations to oxic–anoxic interfaces. Trends Microbiol 13(9):439–448.

3. Kleiner M, et al. (2012) Metaproteomics of a gutless marine worm and its symbiotic microbial community reveal unusual pathways for carbon and energy use. Proc Natl Acad Sci 109(19):E1173–E1182.

4. König S, et al. (2016) Nitrogen fixation in a chemoautotrophic lucinid symbiosis. Nat Microbiol 2:16193.

5. Petersen JM, et al. (2016) Chemosynthetic symbionts of marine invertebrate animals are capable of nitrogen fixation. Nat Microbiol 2:16195.

6. Distel DL, et al. (2017) Discovery of chemoautotrophic symbiosis in the giant shipworm Kuphus polythalamia (Bivalvia: Teredinidae) extends wooden-steps theory. Proc Natl Acad Sci 114(18):E3652–E3658.

7. Dmytrenko O, et al. (2014) The genome of the intracellular bacterium of the coastal bivalve, *Solemya velum*: a blueprint for thriving in and out of symbiosis. BMC Genomics 15(1):924.

8. Gruber-Vodicka HR, et al. (2011) *Paracatenula*, an ancient symbiosis between thiotrophic Alphaproteobacteria and catenulid flatworms. Proc Natl Acad Sci 108(29):12078–12083.

9. Rinke C, et al. (2009) High genetic similarity between two geographically distinct strains of the sulfur-oxidizing symbiont ‘*Candidatus* Thiobios zoothamnicoli.’ FEMS Microbiol Ecol 67(2):229–241.

10. Assié A, et al. (2018) Horizontal acquisition of a patchwork Calvin cycle by symbiotic and free-living Campylobacterota (formerly Epsilonproteobacteria). bioRxiv. doi:10.1101/437616.

11. Suzuki Y, et al. (2005) Novel chemoautotrophic endosymbiosis between a member of the Epsilonproteobacteria and the hydrothermal-vent gastropod *Alviniconcha aff. hessleri* (Gastropoda: Provannidae) from the Indian Ocean. Appl Env Microbiol 71(9):5440–5450.

12. Campbell BJ, Stein JL, Cary SC (2003) Evidence of chemolithoautotrophy in the bacterial community associated with *Alvinella pompejana*, a hydrothermal vent polychaete. Appl Environ Microbiol 69(9):5070–5078.

13. Beinart RA, et al. (2012) Evidence for the role of endosymbionts in regional-scale habitat partitioning by hydrothermal vent symbioses. Proc Natl Acad Sci 109(47):E3241–E3250.

14. Markert S, et al. (2007) Physiological proteomics of the uncultured endosymbiont of Riftia pachyptila. Science 315(5809):247–250.

15. Thiel V, et al. (2012) Widespread occurrence of two carbon fixation pathways in tubeworm endosymbionts: Lessons from hydrothermal vent associated tubeworms from the Mediterranean Sea. Front Microbiol 3:423.

16. Rubin-Blum M, Dubilier N, Kleiner M (2019) Genetic evidence for two carbon fixation pathways (the Calvin-Benson-Bassham cycle and the reverse tricarboxylic acid cycle) in symbiotic and free-living bacteria. mSphere 4(1):e00394–18.

17. Ponsard J, et al. (2013) Inorganic carbon fixation by chemosynthetic ectosymbionts and nutritional transfers to the hydrothermal vent host-shrimp *Rimicaris exoculata*. ISME J 7(1):96–109.

18. Kleiner M, et al. (2015) Use of carbon monoxide and hydrogen by a bacteria–animal symbiosis from seagrass sediments. Environ Microbiol 17(12):5023–5035.

19. Petersen JM, et al. (2011) Hydrogen is an energy source for hydrothermal vent symbioses. Nature 476(7359):176–180.

20. Seah BKB, et al. (2017) Specificity in diversity: single origin of a widespread ciliatebacteria symbiosis. Proc R Soc B Biol Sci 284(1858):20170764.

21. Fenchel T, Finlay BJ (1989) *Kentrophoros*: A mouthless ciliate with a symbiotic kitchen garden. Ophelia 30(2):75–93.

22. Raikov IB (1971) Bactéries épizoiques et mode de nutrition du cilié psammophile *Kentrophoros fistulosum* Fauré-Fremiet (étude au microscope électronique). Protistologica 7(3):365–378.

23. Foissner W (1995) *Kentrophoros* (Ciliophora, Karyorelictea) has oral vestiges: a reinvestigation of *K. fistulosus* (Fauré-Fremiet, 1950) using protargol impregnation. Arch Für Protistenkd 146:165–179.

24. Rodriguez-R LM, Konstantinidis KT (2014) Bypassing cultivation to identify bacterial species. Microbe 9(3):111–8.

25. Tabita FR, et al. (2007) Function, structure, and evolution of the RubisCO-Like Proteins and their RubisCO homologs. Microbiol Mol Biol Rev 71(4):576–599.

26. Dahl C, Friedrich CG (2008) Microbial sulfur metabolism (Springer, Berlin; New York).

27. Ghosh W, Dam B (2009) Biochemistry and molecular biology of lithotrophic sulfur oxidation by taxonomically and ecologically diverse bacteria and archaea. FEMS Microbiol Rev 33(6):999–1043.

28. Pitcher RS, Watmough NJ (2004) The bacterial cytochrome cbb3 oxidases. Biochim Biophys Acta BBA - Bioenerg 1655:388–399.

29. Ducluzeau A-L, Ouchane S, Nitschke W (2008) The cbb3 oxidases are an ancient innovation of the domain Bacteria. Mol Biol Evol 25(6):1158–1166.

30. Kleiner M, Petersen JM, Dubilier N (2012) Convergent and divergent evolution of metabolism in sulfur-oxidizing symbionts and the role of horizontal gene transfer. Curr Opin Microbiol 15(5):621–631.

31. Wood AP, Aurikko JP, Kelly DP (2004) A challenge for 21st century molecular biology and biochemistry: what are the causes of obligate autotrophy and methanotrophy? FEMS Microbiol Rev 28(3):335–352.

32. Fuchs G, Berg IA (2014) Unfamiliar metabolic links in the central carbon metabolism. J Biotechnol 192:314–322.

33. Zarzycki J, Fuchs G (2011) Coassimilation of organic substrates via the autotrophic 3-hydroxypropionate bi-cycle in *Chloroflexus aurantiacus*. Appl Environ Microbiol 77(17):6181–6188.

34. Shih PM, Ward LM, Fischer WW (2017) Evolution of the 3-hydroxypropionate bicycle and recent transfer of anoxygenic photosynthesis into the Chloroflexi. Proc Natl Acad Sci 114(40):10749–10754.

35. Eek KM, Sessions AL, Lies DP (2007) Carbon-isotopic analysis of microbial cells sorted by flow cytometry. Geobiology 5(1):85–95.

36. Kleiner M, et al. (2018) Metaproteomics method to determine carbon sources and assimilation pathways of species in microbial communities. Proc Natl Acad Sci 115(24):E5576–E5584.

37. Boden R, Hutt LP (2018) Chemolithoheterotrophy: Means to higher growth yields from this widespread metabolic trait. Aerobic Utilization of Hydrocarbons, Oils and Lipids, ed Rojo F (Springer International Publishing, Cham), pp 1–25.

38. Kelly DP, Wood AP (2013) The Chemolithotrophic Prokaryotes. The Prokaryotes, eds Rosenberg E, DeLong EF, Lory S, Stackebrandt E, Thompson F (Springer Berlin Heidelberg), pp 275–287.

39. Owen OE, Kalhan SC, Hanson RW (2002) The key role of anaplerosis and cataplerosis for citric acid cycle function. J Biol Chem 277(34):30409–30412.

40. Perez RC, Matin A (1982) Carbon dioxide assimilation by *Thiobacillus novellus* under nutrient-limited mixotrophic conditions. J Bacteriol 150(1):46–51.

41. Roslev P, Larsen MB, Jørgensen D, Hesselsoe M (2004) Use of heterotrophic CO2 assimilation as a measure of metabolic activity in planktonic and sessile bacteria. J Microbiol Methods 59(3):381–393.

42. Hügler M, Sievert SM (2011) Beyond the Calvin cycle: Autotrophic carbon fixation in the ocean. Annu Rev Mar Sci 3(1):261–289.

43. Ott JA, et al. (1991) Tackling the sulfide gradient: a novel strategy involving marine nematodes and chemoautotrophic ectosymbionts. Mar Ecol 12(3):261–279.

44. Conway N, Capuzzo JM, Fry B (1989) The role of endosymbiotic bacteria in the nutrition of *Solemya velum*: evidence from a stable isotope analysis of endosymbionts and host. Limnol Oceanogr 34(1):249–255.

45. Dando PR, Spiro B (1993) Varying nutritional dependence of the thyasirid bivalves *Thyasira sarsi* and *T. equalis* on chemoautotrophic symbiotic bacteria, demonstrated by isotope ratios of tissue carbon and shell carbonate. Mar Ecol-Prog Ser 92:151–151.

46. Hayes JM (2001) Fractionation of carbon and hydrogen isotopes in biosynthetic processes. Rev Mineral Geochem 43(1):225–277.

47. Nunoura T, et al. (2018) A primordial and reversible TCA cycle in a facultatively chemolithoautotrophic thermophile. Science 359(6375):559–563.

48. Mall A, et al. (2018) Reversibility of citrate synthase allows autotrophic growth of a thermophilic bacterium. Science 359(6375):563–567.

49. Ivanovsky RN, Krasilnikova EN, Fal YI (1993) A pathway of the autotrophic CO_2_ fixation in *Chloroflexus aurantiacus*. Arch Microbiol 159(3):257–264.

50. Kuwahara H, et al. (2007) Reduced genome of the thioautotrophic intracellular symbiont in a deep-sea clam, *Calyptogena okutanii*. Curr Biol 17(10):881–886.

51. Newton ILG, et al. (2007) The *Calyptogena magnifica* chemoautotrophic symbiont genome. Science 315(5814):998–1000.

52. Sayavedra L, et al. (2015) Abundant toxin-related genes in the genomes of beneficial symbionts from deep-sea hydrothermal vent mussels. eLife 4:e07966.

53. Yelton AP, et al. (2016) Global genetic capacity for mixotrophy in marine picocyanobacteria. ISME J 10(12):2946–2957.

54. Nelson DC, Williams CA, Farah BA, Shively JM (1988) Occurence and regulation of Calvin cycle enzymes in non-autotrophic *Beggiatoa strains*. Arch Microbiol 151(1):15–19.

55. Childress JJ, Girguis PR (2011) The metabolic demands of endosymbiotic chemoautotrophic metabolism on host physiological capacities. J Exp Biol 214(2):312–325.

56. Ponnudurai R, et al. (2017) Metabolic and physiological interdependencies in the *Bathymodiolus azoricus* symbiosis. ISME J 11(2):463–477.

57. Jørgensen BB (2000) Bacteria and marine biogeochemistry. Marine Geochemistry, eds Schulz HD, Zabel M (Springer, Berlin), pp 173–207. 2nd Ed.

58. Vafeiadou A-M, Materatski P, Adão H, De Troch M, Moens T (2014) Resource utilization and trophic position of nematodes and harpacticoid copepods in and adjacent to *Zostera noltii* beds. Biogeosciences 11(14):4001–4014.

59. Vizzini S, Sara G, Michener RH, Mazzola A (2002) The role and contribution of the seagrass Posidonia oceanica (L.) Delile organic matter for secondary consumers as revealed by carbon and nitrogen stable isotope analysis. Acta Oecologica 23(4):277–285.

60. Cooper LW, DeNiro MJ (1989) Stable carbon isotope variability in the seagrass *Posidonia oceanica*: Evidence for light intensity effects. Mar Ecol Prog Ser 50(3):225–229.

61. McMillan C, Parker PL, Fry B (1980) 13C/12C ratios in seagrasses. Aquat Bot 9:237–249.

62. Heuer V, et al. (2006) Online δ13C analysis of volatile fatty acids in sediment/porewater systems by liquid chromatography-isotope ratio mass spectrometry. Limnol Oceanogr Methods 4(10):346–357.

63. Høgslund S, et al. (2009) Physiology and behaviour of marine Thioploca. ISME J 3(6):647–657.

64. Winkel M, et al. (2016) Single-cell sequencing of *Thiomargarita reveals* genomic flexibility for adaptation to dynamic redox conditions. Front Microbiol 7:964.

65. Seviour RJ, McIlroy S (2008) The microbiology of phosphorus removal in activated sludge processes-the current state of play. J Microbiol 46(2):115–124.

66. Schulz HN, Schulz HD (2005) Large sulfur bacteria and the formation of phosphorite. Science 307(5708):416–418.

67. Medlin L, Elwood HJ, Stickel S, Sogin ML (1988) The characterization of enzymatically amplified eukaryotic 16S-like rRNA-coding regions. Gene 71:491–499.

68. Bankevich A, et al. (2012) SPAdes: a new genome assembly algorithm and its applications to single-cell sequencing. J Comput Biol 19(5):455–477.

69. Peng Y, Leung HCM, Yiu SM, Chin FYL (2012) IDBA-UD: a de novo assembler for single-cell and metagenomic sequencing data with highly uneven depth. Bioinformatics 28(11):1420–1428.

70. Wu M, Scott AJ (2012) Phylogenomic analysis of bacterial and archaeal sequences with AMPHORA2. Bioinformatics 28(7):1033–1034.

71. Wang Z, Wu M (2013) A phylum-level bacterial phylogenetic marker database. Mol Biol Evol 30(6):1258–1262.

72. Quast C, et al. (2013) The SILVA ribosomal RNA gene database project: improved data processing and web-based tools. Nucleic Acids Res 41(D1):D590–D596.

73. Edgar RC (2010) Search and clustering orders of magnitude faster than BLAST. Bioinformatics 26(19):2460–2461.

74. Albertsen M, et al. (2013) Genome sequences of rare, uncultured bacteria obtained by differential coverage binning of multiple metagenomes. Nat Biotechnol 31(6):533–538.

75. Seah BKB, Gruber-Vodicka HR (2015) gbtools: Interactive visualization of metagenome bins in R. Front Microbiol 6:1451.

76. Gurevich A, Saveliev V, Vyahhi N, Tesler G (2013) QUAST: quality assessment tool for genome assemblies. Bioinformatics 29(8):1072–1075.

77. Parks DH, Imelfort M, Skennerton CT, Hugenholtz P, Tyson GW (2015) CheckM: assessing the quality of microbial genomes recovered from isolates, single cells, and metagenomes. Genome Res 25:1043–1055.

78. Richter M, Rosselló-Móra R (2009) Shifting the genomic gold standard for the prokaryotic species definition. Proc Natl Acad Sci 106(45):19126–19131.

79. Markowitz VM, et al. (2014) IMG/M 4 version of the integrated metagenome comparative analysis system. Nucleic Acids Res 42(D1):D568–D573.

80. Karp PD, Latendresse M, Caspi R (2011) The Pathway Tools pathway prediction algorithm. Stand Genomic Sci 5(3):424–429.

81. Karp PD, et al. (2016) Pathway Tools version 19.0 update: software for pathway/genome informatics and systems biology. Brief Bioinform 17(5):877–890.

82. Kanehisa M, Furumichi M, Tanabe M, Sato Y, Morishima K (2017) KEGG: new perspectives on genomes, pathways, diseases and drugs. Nucleic Acids Res 45(D1):D353–D361.

83. Camacho C, et al. (2009) BLAST+: architecture and applications. BMC Bioinformatics 10(1):421.

84. van Dongen S, Abreu-Goodger C (2012) Using MCL to extract clusters from networks. Bacterial Molecular Networks, eds van Helden J, Toussaint A, Thieffry D (Springer New York, New York, NY), pp 281–295.

85. Li L, Stoeckert CJ, Roos DS (2003) OrthoMCL: identification of ortholog groups for eukaryotic genomes. Genome Res 13(9):2178–2189.

86. Wattam AR, et al. (2014) PATRIC, the bacterial bioinformatics database and analysis resource. Nucleic Acids Res 42(D1):D581–D591.

87. Liao Y, Smyth GK, Shi W (2014) featureCounts: an efficient general purpose program for assigning sequence reads to genomic features. Bioinformatics 30(7):923–930.

88. Bar-Even A, Noor E, Milo R (2012) A survey of carbon fixation pathways through a quantitative lens. J Exp Bot 63(6):2325–2342.

89. Berg IA (2011) Ecological aspects of the distribution of different autotrophic CO2 fixation pathways. Appl Environ Microbiol 77(6):1925–1936.

90. Fuchs G (2011) Alternative pathways of carbon dioxide fixation: Insights into the early evolution of life? Annu Rev Microbiol 65(1):631–658.

91. Magrane M, Consortium U (2011) UniProt Knowledgebase: a hub of integrated protein data. Database 2011:bar009.

92. Buchfink B, Xie C, Huson DH (2014) Fast and sensitive protein alignment using DIAMOND. Nat Methods 12(1):59–60.

93. Saier MH, Reddy VS, Tamang DG, Vastermark A (2014) The Transporter Classification Database. Nucleic Acids Res 42(D1):D251–D258.

94. Krogh A, Larsson B, von Heijne G, Sonnhammer EL. (2001) Predicting transmembrane protein topology with a hidden Markov model: application to complete genomes. J Mol Biol 305(3):567–580.

95. Price MN, Dehal PS, Arkin AP (2010) FastTree 2–approximately maximum-likelihood trees for large alignments. PloS One 5(3):e9490.

96. Edgar RC (2004) MUSCLE: multiple sequence alignment with high accuracy and high throughput. Nucleic Acids Res 32(5):1792–1797.

97. Wiśniewski JR, Zougman A, Nagaraj N, Mann M (2009) Universal sample preparation method for proteome analysis. Nat Methods 6(5):359–362.

98. Hamann E, et al. (2016) Environmental Breviatea harbour mutualistic Arcobacter epibionts. Nature 534(7606):254–258.

99. Kleiner M, et al. (2017) Assessing species biomass contributions in microbial communities via metaproteomics. Nat Commun 8(1):1558.

100. Fu L, Niu B, Zhu Z, Wu S, Li W (2012) CD-HIT: accelerated for clustering the nextgeneration sequencing data. Bioinformatics 28(23):3150–3152.

101. Chambers MC, et al. (2012) A cross-platform toolkit for mass spectrometry and proteomics. Nat Biotechnol 30(10):918–920.

