## Supplementary Information for "Sulfur-oxidizing symbionts without canonical genes for autotrophic CO_2_ fixation"

to accompany

### Supplementary Materials and Methods

#### *Metabolite extraction and identification*

*Kentrophoros* sp. H was collected on Elba in 2014 for metabolomics (Supplementary Table 7). Samples were fixed in 1 mL cold methanol (HPLC-grade, Sigma-Aldrich) and stored at -20°C until use. Ribitol (40 µL, 200 mg $L^{-1}$, aqueous) was added as internal standard. For metabolite extraction, each sample was resuspended by vortexing, transferred to a bead-beating vial (FastPrep Lysing Matrix B, MP Biomedicals), and disrupted (4 $ms^{-1}$, 40 s). The vial was centrifuged (16000 rcf, 2 min) and supernatant transferred to a new tube. AMW mixture (1 mL acetonitrile/methanol/water in 2:2:1 ratio) was added to the bead-beating vial, which was disrupted and centrifuged again. Supernatant containing metabolites was pooled, and evaporated to dryness under vacuum (Concentrator Plus, Eppendorf, V-AL mode, 30°C, 4 h).

Dried samples were derivatized with methoxyamine hydrochloride (MeOX) dissolved in pyridine and N,O-bis(trimethylsilyl)trifluoroacetamide with 1% trimethylchlorosilane (BSTFA + 1% TMCS). For each sample, MeOX (80 µL, 20 mg $mL^{-1}$) was added, briefly vortexed, and then heated with shaking (37 °C, 1200 rpm, 90 min). The pyridine was evaporated under a stream of $N_2$ gas until samples were dry (≥1 h). 80 µL of BSTFA (Chromatographie Service) was added, vortexed, and heated with shaking (37 °C, 1350 rpm, 30 min). Samples were briefly centrifuged down and transferred to glass vials for GC-MS (Insert G27, spring S27, Mikro-KH-Vial G1; Chromatographie Service). GC-MS analysis was performed on a 7890B GC system (Agilent Technologies) coupled to a 5977A MSD (Agilent). He was used as carrier gas at constant flow of 1 mL $min^{-1}$. The temperature program was 60 °C (2min), increase to 300 °C at 10 °C $min^{-1}$, hold at 325 °C (7 min). The quadrupole MS was operated in electron ionization mode at 70 eV, with scanning range set to 50-600 m/z.

GC-MS data were screened with AMDIS software for known metabolites. Mass spectra were deconvoluted with the AMDIS algorithm ("simple" mode), and searched against an in-house database. Quantification was performed with Agilent Quantiative Analysis software. A quantitation method was created from acquired scan data with the built-in deconvolution algorithm, using default settings except that m/z value 73 was excluded as a quantifier because it corresponds to a derivitization product (trimethylsilyl group). Predicted compounds were assigned identities based on matches to the NIST database and/or in-house database. The method was applied to all samples and blanks as a batch, with default values except GC retention time window of 0.10 min. Quantifier and qualifier peaks were manually curated to correct misassigned peaks. Compounds present in both blanks and samples in similar quantities were excluded.

#### *Screening for hypothetical autotrophic pathway genes in bacterial genomes*

Five (meta)genomes known to have an incomplete 3HPB pathway were screened for genes that could hypothetically allow for autotrophic $CO_2$ fixation, using the Gene Profile tool in IMG/ER (unidirectional sequence similarities, Blastp cutoffs at 10% identity, E-value < 0.1): *Ca*. Thiosymbion from *Olavius algarvensis*, *Ca*. Accumulibacter, and the Pink Berry consortium metagenome). Sequences from two Kentron genomes were used as queries. Nine other genomes belonging to other thiotrophic symbiotic bacteria were also screened with the same criteria. Accession numbers are given in Supplementary File 1.

##### *Screening for lithoheterotrophic metabolism in bacterial genomes*

Genomes available on the IMG/ER platform were screened for genes related to thiotrophic carbon metabolism and carbon fixation (CBB and rTCA cycles), using the following KEGG Orthology (KO) numbers. For sulfur oxidation: K11180, K11181, K17230, K17229, K17222, K17224, K17225, K17223, K17226, K17227, K17218. For CBB and rTCA cycles: K15230, K15231, K15234, K01601, K01602. For each KO term, the list of genomes containing a gene annotated with that KO number was retrieved, filtered to domain Bacteria, and including all genomes "All Finished, Permanent Draft, and Draft".

Both reductive and oxidative (reverse) DsrAB are included under the same KO numbers. To distinguish between the two, the DsrAB amino acid sequences were downloaded from IMG, and aligned by Blastp against a database of DsrAB sequences that have been classified into oxidative and reductive types (1). The best hit was used to annotate the query sequences. However, a strain of *Desulfovibrio alkaliphilus* with reductive-type DsrAB (based on sequence homology) has recently been shown to be able to run the pathway in the oxidative direction, so the pathway may be more flexible than previously thought (2).

Incomplete genomes may give a false positive result of lithoheterotrophy. For the set of candidate lithoheterotrophs with the composite rDsr-Sox pathway, completeness was estimated with the CheckM pipeline (lineage workflow, reduced tree), and genomes with <75% estimated completeness were excluded.

#### Supplementary Results and Discussion

##### *Metabolites detected in* Kentrophoros

*Kentrophoros* sp. H was used for metabolite profiling as it was the largest morphospecies known to us, and could be reliably collected from one site. The predominant metabolite detected was the disaccharide trehalose. Sucrose was also detected, but in smaller quantities. Other possible metabolites detected were also found in the blanks and hence disregarded. Trehalose is a disaccharide of glucose that is phylogenetically widespread, found in both eukaryotes and prokaryotes (3). Only two Kentron phylotypes have the potential for trehalose synthesis, via trehalose synthase, but none have pathways for trehalose breakdown (e.g. trehalase) nor PTS-type sugar uptake transporters. Therefore, trehalose is likely produced and stored by the host ciliates, where they could function as either an energy store or an osmoprotectant.

#### *Reactions that potentially allow for autotrophic $CO_2$ fixation*

Although no known pathways for autotrophic $CO_2$ fixation were predicted in Kentron genomes, several enzymes predicted in Kentron genomes can potentially catalyze a network of reactions that would allow autotrophic $CO_2$ fixation, by combining reactions of the 3-hydroxypropionate bi-cycle (3HPB) (4) and a hypothetical pathway that was previously proposed for *Chloroflexus* (5). The original version of the proposed Ivanovsky pathway by itself would have presented a metabolic dead end, as its net product is glyoxylate, and the only other predicted enzyme in Kentron that metabolizes glyoxylate, malate synthase, effectively reverses the last step of the Ivanovsky pathway. Other reactions that could convert glyoxylate to downstream metabolites, such as tartronate semialdehyde synthase, or isocitrate lyase (part of the glyoxylate shunt), were not predicted for Kentron. However, by allowing the interconversion between the reactant pairs acetyl-CoA/pyruvate and propionyl-CoA/glyoxylate via C5 intermediates in reactions that were previously thought to be unique to the 3HPB, it is possible to form a closed cycle whose net product is pyruvate (Supplementary Table 8).

Some of these reactions would be catalyzed by alternative enzymes compared to the versions originally presented in (5) or (6): PEP carboxykinase (GDP) instead of PEP carboxylase, succinyl-CoA:malate CoA transferase instead of malyl-CoA synthetase, and pyruvate phosphate dikinase instead of pyruvate water kinase. Genes for all the components of the hypothetical pathway were expressed in the transcriptomes sequenced (Supplementary File 1). The putative methylmalonyl-CoA epimerases were originally annotated as lactoylglutathione lyase in IMG, but as the sequences are relatively short (ca. 150 a.a.), they were re-evaluated as likely methylmalonyl-CoA epimerases because their genes were frequently adjacent to those for acyl-CoA carboxyltransferases, and an InterPro search found signatures such as the VOC domain (InterPro IPR037523) and Glyoxalase_4 domain (Pfam PF13669) which are also characteristic of methylmalonyl-CoA epimerase, but not the signatures characteristic of lactoylglutathione lyase (e.g. IPR019883 and IPR004361).

#### Thermodynamic favorability of the proposed reactions

The overall net reaction is exergonic (-60.9 kJ/mol), but some individual steps are endergonic and could be significant barriers. The most positive $\Delta_r G'^m$ values are for pyruvate synthase (18.7 kJ/mol) and pyruvate phosphate dikinase (19.6 kJ/mol), whereas carboxylation of PEP requires 4.3 kJ/mol. However, for the phosphorylation of pyruvate, the by-product pyrophosphate is very favorably hydrolyzed to two orthophosphate groups, and the net reaction is exergonic (-13.3 kJ/mol) and would be favored if they were coupled. The hydrolysis of pyrophosphate can be coupled to energy conservation by $Na^+/H^+$-translocating pyrophosphatase, which is predicted in all Kentron genomes.

The pyruvate synthase reaction can also be more favorable if the ratio of reduced:oxidized ferredoxin (assuming reduction potential of -418 +/- 60 mV) in the cell is on the order of 100-fold or greater (Supplementary Table 9). The value of +18.7 kJ/mol is for 1 mM concentrations of both reduced and oxidized species. At a ratio of 100:1, the reaction could be exergonic (although the uncertainty is +/-13.2 kJ/mol). Such an “over-reduced” state in the cell could be maintained by the Rnf-type $Na^+$-translocating oxidoreductases, which can reduce ferredoxin with NADH using energy from a $Na^+$ membrane gradient. These are predicted in Kentron genomes, and are relatively common among facultatively anaerobic bacteria (7).

In kinetic terms, however, pyruvate synthase is slow (specific activity < 0.1 to 2.3 $\mu mol\ min^{-1}\ mg^{-1}$) and has low substrate specificity ($K_M$ 2 to 48 mM) in organisms where these parameters have been measured; in comparison, RuBisCO has a specific activity of 2 to 4 $\mu mol\ min^{-1}\ mg^{-1}$ (cited in (6)). This means that pyruvate synthase would have to be highly expressed, at a level comparable to RuBisCO in CBB-cycle organisms, for it to be an effective autotrophic carboxylase. In Kentron H, pyruvate synthase was among the top 5% in expression level (Figure 4), but this was only about 6% of the expression level of the top-expressed gene (a predicted phasin). In comparison, RuBisCO is typically the most-expressed, or among the most-expressed genes in thiotroph transcriptomes (8, 9). Nonetheless, as the organisms had to be extracted from their natural sediment habitat before being fixed, it is likely that the gene expression levels do not reflect their *in situ* metabolic states.

#### Role of these reactions in Kentron and other bacteria

Other bacterial genomes that encode enzymes for an incomplete 3HPB, namely *Ca*. Thiosymbion and *Ca*. Accumulibacter, also encoded genes for other enzymes of the hypothetical pathway (Supplementary File 1). *Ca*. Thiosymbion are is a thiotrophic symbionts like Kentron, but they already possess a functioning CBB cycle (10). In comparison, *Ca*. Accumulibacter are chemoorganotrophs from wastewater treatment plants, and it is unclear why they would possess an autotrophic pathway (some *Ca*. Accumulibacter also encode a CBB cycle).

Like the complete 3HPB in *Chloroflexus* (11), the hypothetical pathway would also allow the co-assimilation of organic substrates, such as succinate, malate, and propionate. Given that (i) such organic acids are predicted substrates of the uptake transporters encoded in Kentron genomes and are also common in coastal sediments, (ii) the reactions are also predicted in a thiotrophic symbiont that uses the CBB cycle for autotrophy, and (iii) the 3HPB is used mixotrophically in *Chloroflexus*, it is likely that the usual nutritional mode of Kentron is lithoheterotrophic, or at most mixotrophic. If $CO_2$ fixation only occurs with organic co-assimilation, it would then be indistinguishable from heterotrophic $CO_2$ fixation, especially as the carboxylases involved are also typical for carboxylation in heterotrophs.

#### *Oxidation/reduction value of substrates and biomass*

The oxidation/reduction (O/R) value can be used as a measure of how oxidized or reduced a substrate is relative to biomass, and hence whether additional reducing equivalents are required for its assimilation (12). For a molecular formula where the ratio of H:O is $x$:$y$, the oxidation/reduction level $r = (2y - x)/2$. The empirical formula $CH_{1.77}O_{0.49}N_{0.24}$ of biomass for *Escherichia coli* was used (13), which is close to empirical formulas for other bacteria, e.g. the purple non-sulfur bacterium *Rhodopseudomonas palustris* $CH_{1.8}O_{0.38}N_{0.18}$ (14). O/R values for potential substrates and storage compounds of Kentron are shown in Supplementary Table 4. Assimilation of malate, succinate, acetate and the mobilization of glycogen for biosynthesis would require additional reducing equivalents, whereas polyhydroxybutyrate (a form of PHA) could serve as a store of reducing equivalents.

##### *Occurrence of lithoheterotrophic metabolism in thiotroph genomes*

A total of 1407 thiotrophic bacterial genomes in the IMG/ER database (excluding Kentron) were predicted to encode lithoheterotrophic metabolism, based on a screening using key genes for sulfur oxidation and autotrophy (CBB and rTCA cycles). Genomes that encode the rDsr/Sox pathway but lack either CBB or rTCA cycles are uncommon – only seven genomes were found (Supplementary Table 10). One of these, *Magnetococcus marinus* MC-1 was a false negative, as it has been reported to use the rTCA cycle (15), but it encodes a variant of the ATP citrate lyase that was not annotated with a KO number by the IMG pipeline. The others were *Ruegeria marina* CGMCC 1.9108, *Thiothrix flexilis* DSM 14609, and four genome bins from environmental metagenomes: *Thioalkalivibrio* spp. HK1 and TsSOB (both associated with sponges), REDSEA-S14_B17, and REDSEA-S15_B12. Most genomes that encode the rDsr/Sox pathway have either the CBB cycle (89 genomes) or both (8). In contrast, those that encode the Sox pathway only are roughly as likely to have at least one autotrophic pathway (642) as not (661).

The Sox pathway alone allows oxidation of reduced sulfur in the form of thiosulfate. Additional enzymes (Sqr and FccAB) would also allow oxidation of sulfide. However, bacteria that can store elemental sulfur or polysulfide as cellular inclusions typically have the rDsr/Sox pathway, because the Dsr components are required to mobilize the stored sulfur (16). We hypothesize that the rDsr/Sox pathway is more often associated with autotrophic pathways because $CO_2$ fixation can secondarily serve as an additional electron sink under reducing conditions, when other electron acceptors such as oxygen or nitrate are unavailable.

#### References for Supplement


1. Müller AL, Kjeldsen KU, Rattei T, Pester M, Loy A (2015) Phylogenetic and environmental diversity of DsrAB-type dissimilatory (bi)sulfite reductases. *ISME J* 9:1152–1165.

2. Thorup C, Schramm A, Findlay AJ, Finster KW, Schreiber L (2017) Disguised as a sulfate reducer: Growth of the deltaproteobacterium *Desulfurivibrio alkaliphilus* by sulfide oxidation with nitrate. *mBio* 8(4):e00671-17.

3. Elbein AD, Pan YT, Pastuszak I, Carroll D (2003) New insights on trehalose: a multifunctional molecule. *Glycobiology* 13(4):17R-27R.

4. Zarzycki J, Brecht V, Müller M, Fuchs G (2009) Identifying the missing steps of the autotrophic 3-hydroxypropionate $CO_2$ fixation cycle in *Chloroflexus aurantiacus*. *Proc Natl Acad Sci* 106(50):21317–21322.

5. Ivanovsky RN, Krasilnikova EN, Fal YI (1993) A pathway of the autotrophic $CO_2$ fixation in *Chloroflexus aurantiacus*. *Arch Microbiol* 159(3):257–264.

6. Bar-Even A, Noor E, Milo R (2012) A survey of carbon fixation pathways through a quantitative lens. *J Exp Bot* 63(6):2325–2342.

7. Biegel E, Schmidt S, González JM, Müller V (2011) Biochemistry, evolution and physiological function of the Rnf complex, a novel ion-motive electron transport complex in prokaryotes. *Cell Mol Life Sci* 68(4):613–634.

8. Seston SL, et al. (2016) Metatranscriptional response of chemoautotrophic *Ifremeria nautilei* endosymbionts to differing sulfur regimes. *Front Microbiol* 7:1074.

9. Stewart FJ, Dmytrenko O, DeLong EF, Cavanaugh CM (2011) Metatranscriptomic analysis of sulfur oxidation genes in the endosymbiont of *Solemya velum*. *Front Microbiol* 2:134.

10. Kleiner M, et al. (2012) Metaproteomics of a gutless marine worm and its symbiotic microbial community reveal unusual pathways for carbon and energy use. *Proc Natl Acad Sci* 109(19):E1173–E1182.

11. Zarzycki J, Fuchs G (2011) Coassimilation of organic substrates via the autotrophic 3-hydroxypropionate bi-cycle in *Chloroflexus aurantiacus*. *Appl Environ Microbiol* 77(17):6181–6188.

12. Gottschalk G (1986) *Bacterial Metabolism* (Springer New York, New York, NY) doi:10.1007/978-1-4612-1072-6.

13. Grosz R, Stephanopoulos G (1983) Statistical mechanical estimation of the free energy of formation of *E. coli* biomass for use with macroscopic bioreactor balances. *Biotechnol Bioeng* 25(9):2149–2163.

14. McKinlay JB, Harwood CS (2010) Carbon dioxide fixation as a central redox cofactor recycling mechanism in bacteria. *Proc Natl Acad Sci* 107(26):11669–11675.

15. Williams TJ, Zhang CL, Scott JH, Bazylinski DA (2006) Evidence for autotrophy via the reverse tricarboxylic acid cycle in the marine magnetotactic coccus strain MC-1. *Appl Environ Microbiol* 72(2):1322–1329.

16. Ghosh W, Dam B (2009) Biochemistry and molecular biology of lithotrophic sulfur oxidation by taxonomically and ecologically diverse bacteria and archaea. *FEMS Microbiol Rev* 33(6):999–1043.

17. Flamholz A, Noor E, Bar-Even A, Milo R (2012) eQuilibrator--the biochemical thermodynamics calculator. *Nucleic Acids Res* 40(D1):D770–D775.

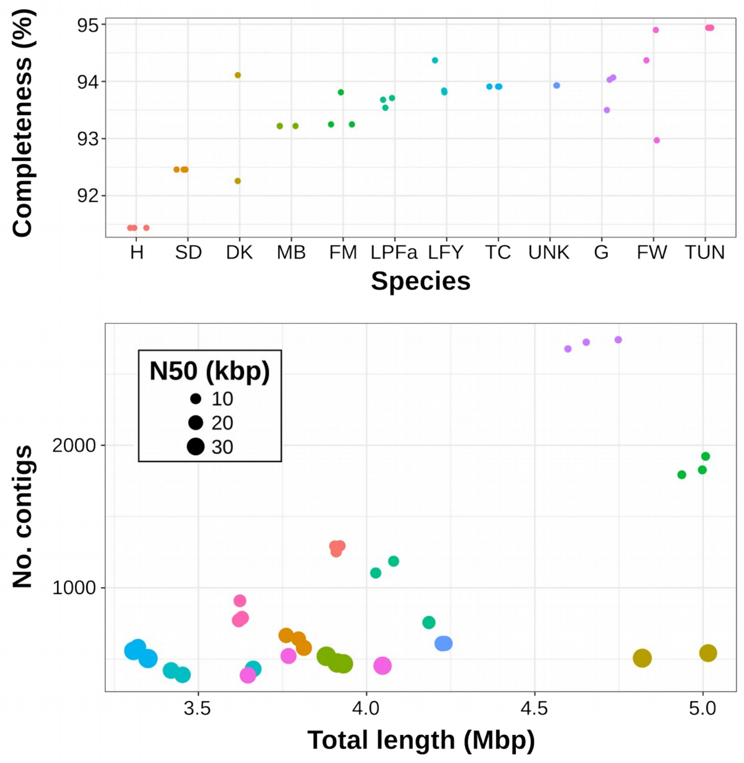


**Supplementary Figure 1**. Summary statistics for Kentron genome assemblies. *Above*: Genome completeness per species, as assessed with gammaproteobacterial conserved marker genes by CheckM. *Below:* Number of contigs vs. total length (Mbp) of Kentron genome bins; plot symbol areas scaled by N50 (kbp). Larger genomes are usually more fragmented (more contigs, lower N50), with the exception of Kentron sp. DK.

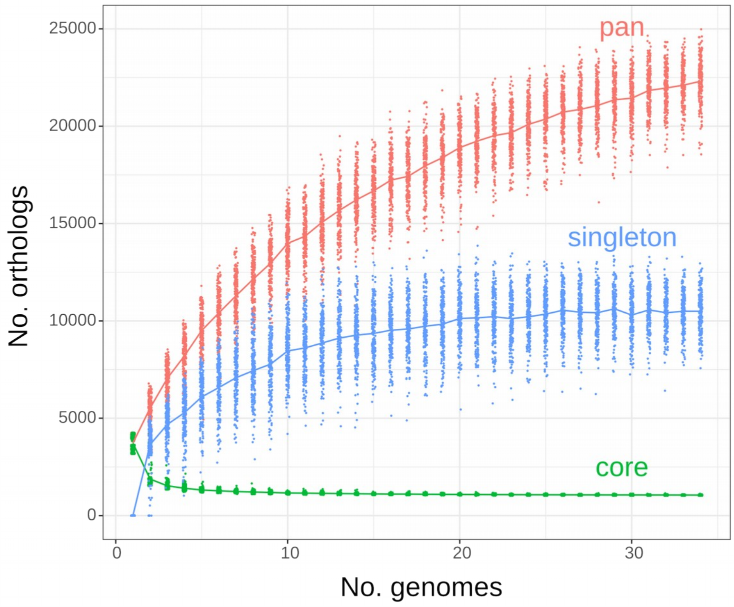


255 **Supplementary Figure 2**. Accumulation curves for pan and core genomes (protein-coding
256 genes only) of the Kentron clade, with uncertainty estimated by resampling (200 times, with
257 replacement). “Singleton” genes were not included in an ortholog cluster.

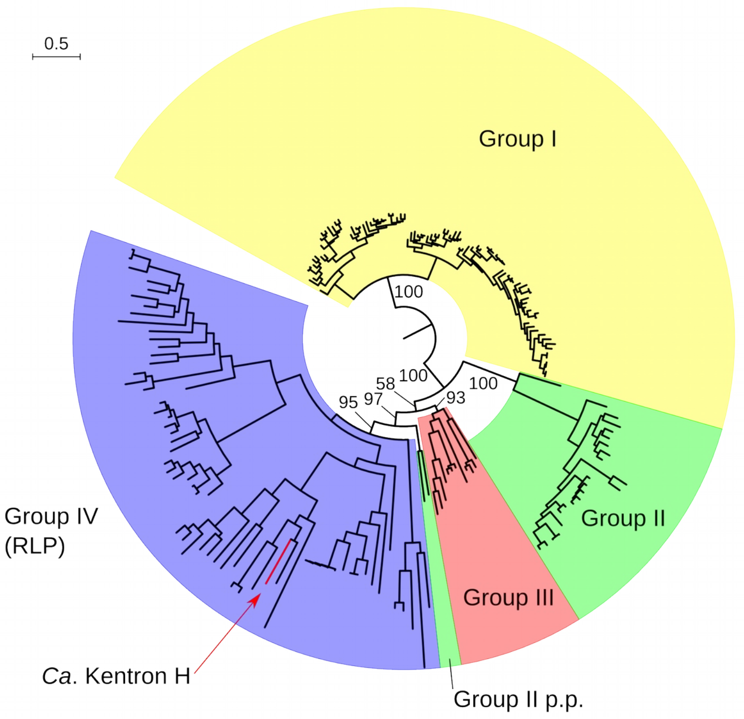


**Supplementary Figure 3.** Maximum-likelihood phylogeny of RuBisCO proteins including RuBisCO-like protein from Kentron phylotype H (red arrow).

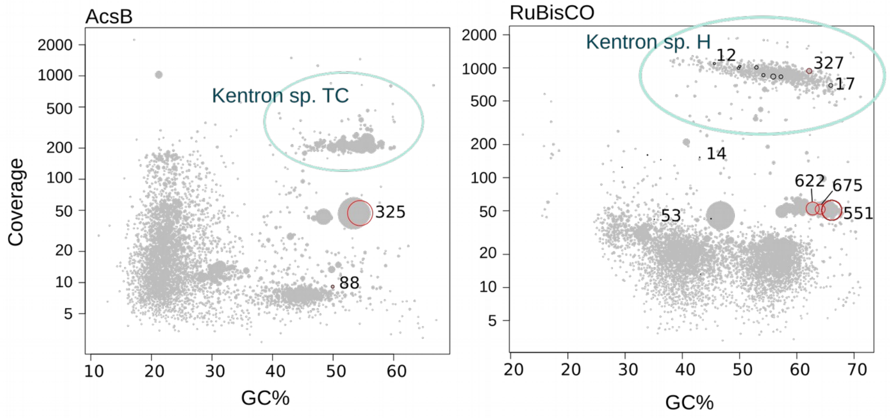


**Supplementary Figure 4**. Selected results of re-mapping reads with hits to canonical autotrophy enzymes to the metagenomic assemblies, for AcsB vs. *Kentrophoros* sp. TC (*left*) and RuBisCO vs. *Kentrophoros* sp. H (*right*). Metagenome assemblies are depicted as coverage-GC% plots, where each grey circle represents a contig, with its size scaled to the contig length. Contigs with remapping hits have solid outlines, with numbers of remapping reads indicated when >10. The majority of reads with alignments to canonical autotrophy enzymes map to scaffolds outside the Kentron symbiont genome bin (circled in blue) and could be attributed to other bacteria in the metagenomes, except for RuBisCO in Kentron sp. H, due to the RuBisCO-like protein encoded in that phylotype.

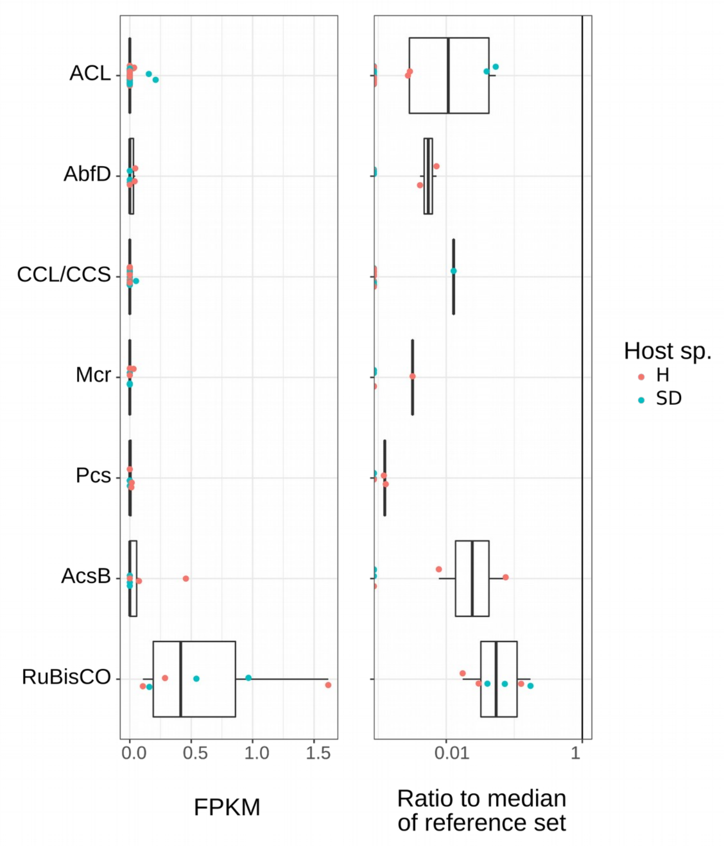


**Supplementary Figure 5**. Read coverage in *Kentrophoros* metatranscriptomes for key enzymes of autotrophic $CO_2$-fixation pathways, expressed as FPKM values (*left*) and as a fraction of the median coverage of a reference set of proteins (*right*). Each point represents a separate metatranscriptome library, colored by *Kentrophoros* host morphospecies. Abbreviations as in Figure 3.

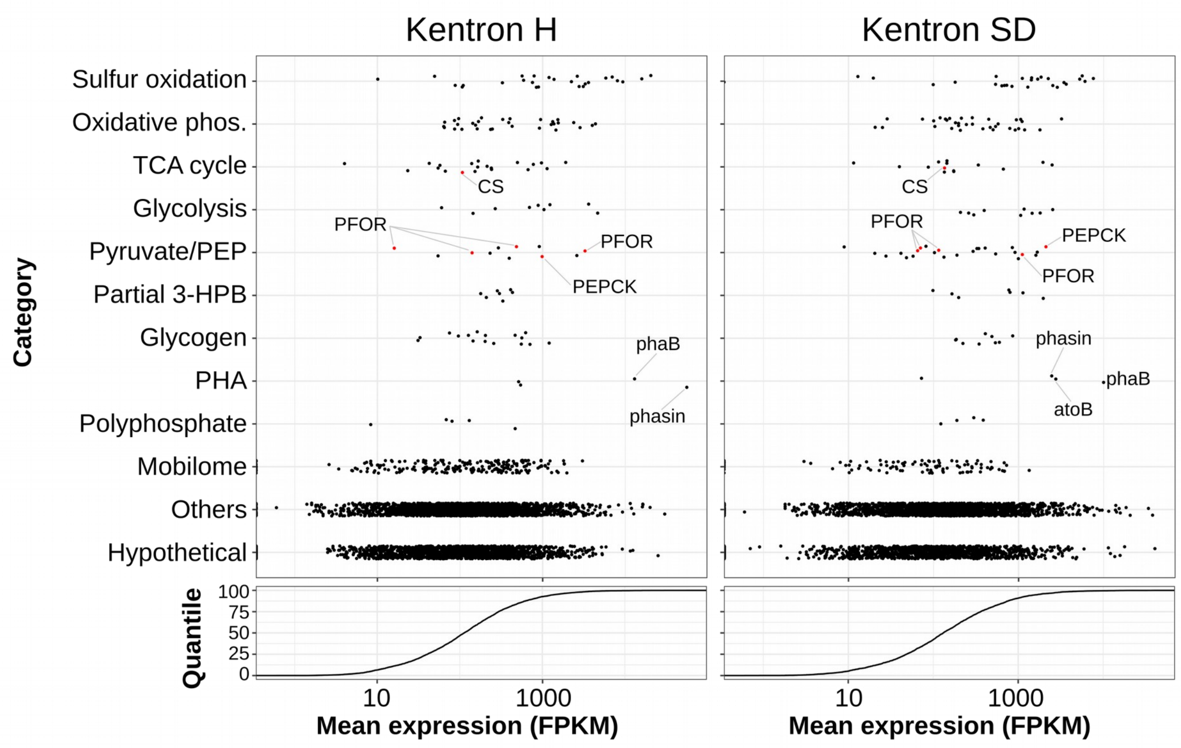


**Supplementary Figure 6**. Expression of genes in selected functional categories, for two Kentron phylotypes (mean of 3 samples per phylotype, in FPKM). Cumulative distribution curves for expression per gene are shown below each dot-plot. Selected genes discussed in text are labeled; carboyxlation reactions involving pyruvate or phosphoenolpyruvate (PEP) are marked in red. *Abbreviations*: atoB, acetyl-CoA acetyltransferase; CS, citrate synthase; PFOR, pyruvate-ferredoxin oxidoreductase (= pyruvate synthase); PEPCK, phosphenolpyruvate carboxykinase; phbB, acetoacetyl-CoA reductase.

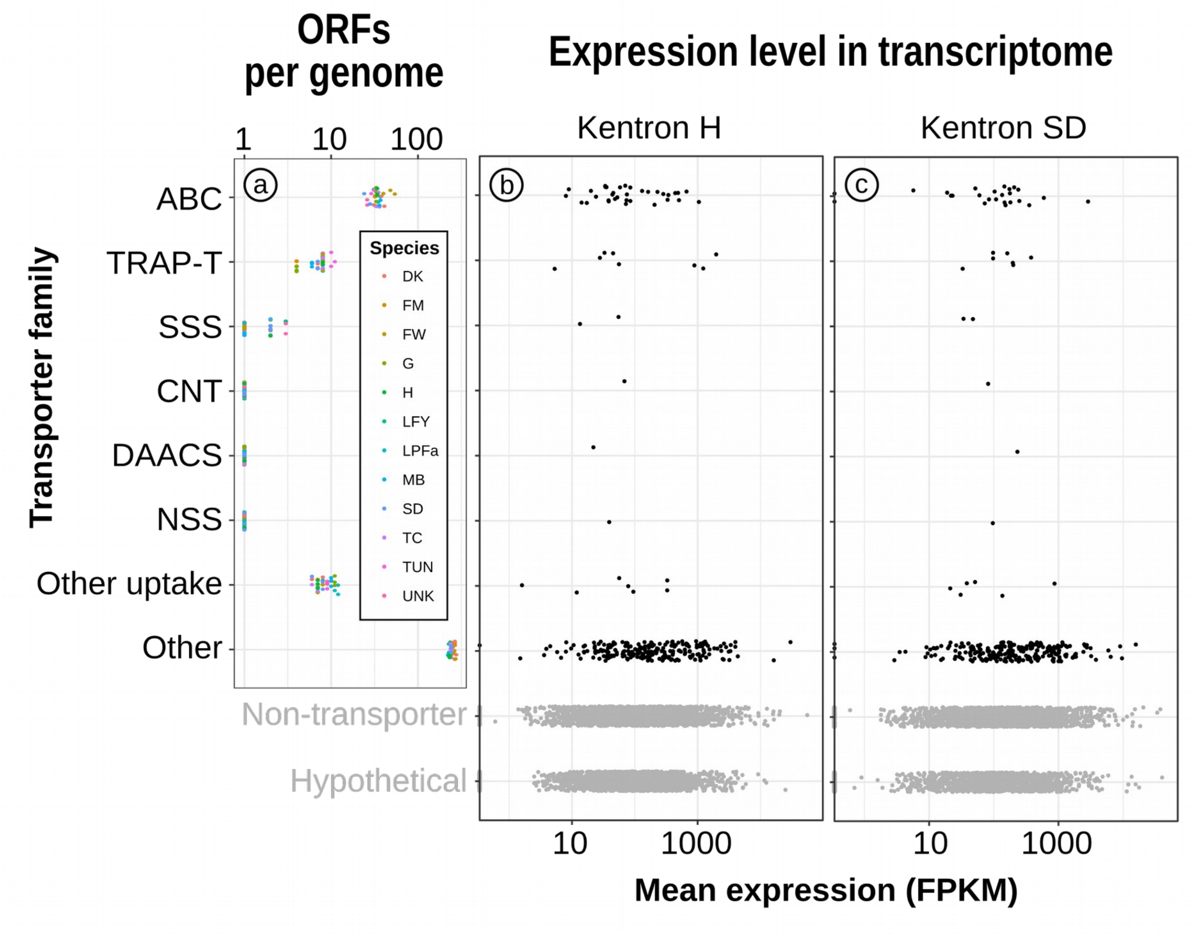


**Supplementary Figure 7**. Uptake-related transporter families found in all Kentron genomes. (a) No. of open reading frames per genome with matches to Transporter Classification families (Blastp E-value <10-5, >30% amino acid identity, >70% query coverage). (b) Mean expression (FPKM) in three transcriptomes of Kentron H. (c) Mean expression (FPKM) in three transcriptomes of Kentron SD.

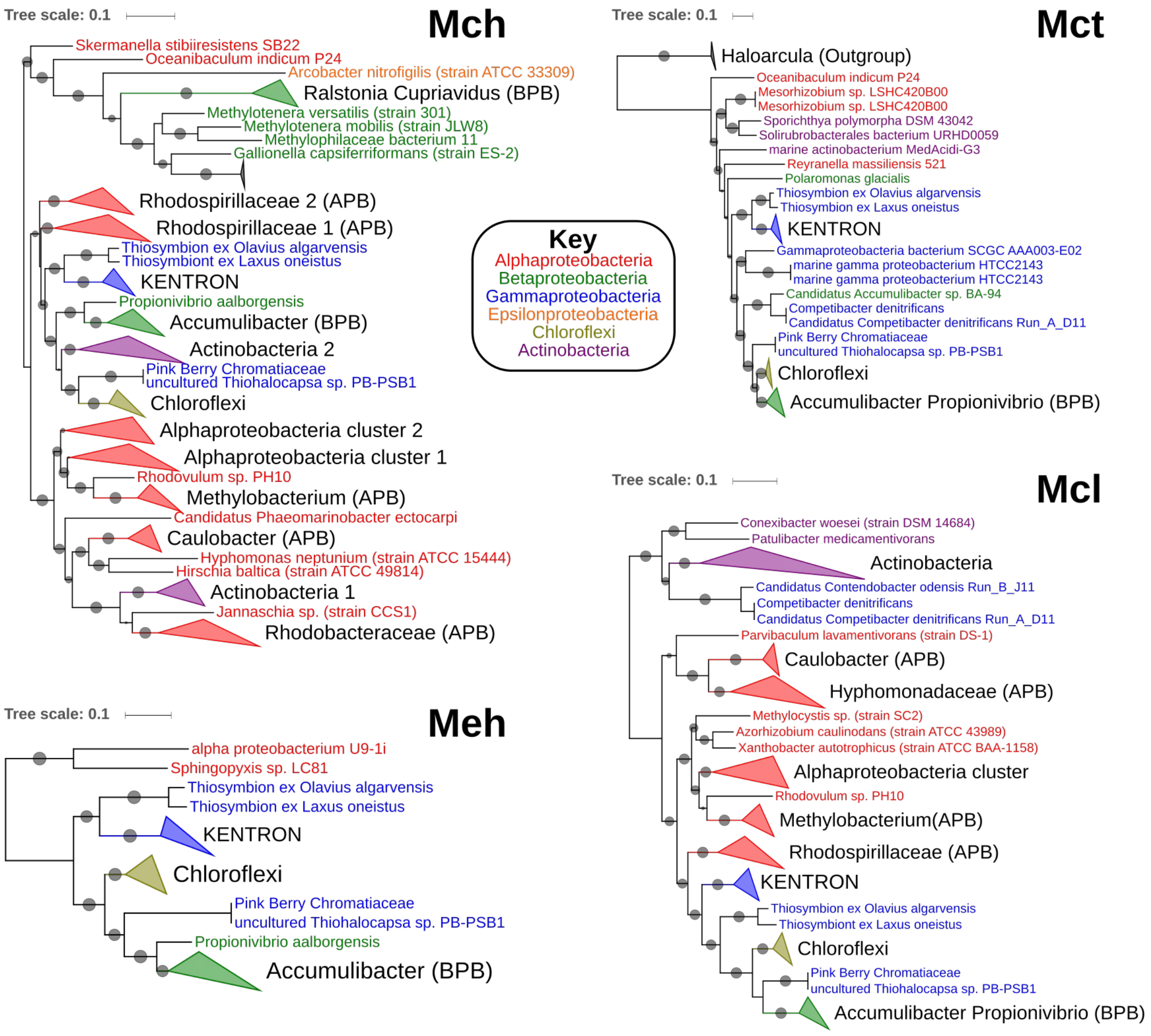


**Supplementary Figure 8**. Phylogenetic trees of genes of 3-hydroxypropionate bi-cycle found in Kentron, with homologs from other bacteria. *Abbreviations*: Mch, mesaconyl-C1-CoA hydratase; Mcl, malyl-CoA/beta-methylmalyl-CoA/citramalyl-CoA (MMC) lyase; Mct, mesaconyl-CoA C1-C4 CoA transferase; Meh, mesaconyl-C4-CoA hydratase.

**Supplementary Table 1**. Collection localities and dates for *Kentrophoros* metagenomics (MG) and transcriptomics (T) samples.

| Purpose | Library | Sample no. | BioSample accession | Locality | Latitude | Longitude | Date | Host |
|---|---|---|---|---|---|---|---|---|
| MG | 772A | SA2B15 | SAMEA104679830 | Italy; Elba; Sant' Andrea | 42.808561 °N | 10.142275 °E | 2013-08-11 | H |
| MG | 772B | SA2B20 | SAMEA104679831 | Italy; Elba; Sant' Andrea | 42.808561 °N | 10.142275 °E | 2013-08-11 | H |
| MG | 772C | SA2B12 | SAMEA104679832 | Italy; Elba; Sant' Andrea | 42.808561 °N | 10.142275 °E | 2013-08-11 | H |
| MG | 772D | S127 | SAMEA104679833 | Italy; Elba; Fetovaia | 42.7313 °N | 10.1534 °E | 2013-07-02 | SD |
| MG | 772E | S1320 | SAMEA104679834 | Italy; Elba; Fetovaia | 42.7313 °N | 10.1534 °E | 2013-07-02 | SD |
| MG | 772F | S1321 | SAMEA104679835 | Italy; Elba; Fetovaia | 42.7313 °N | 10.1534 °E | 2013-07-02 | SD |
| MG | 1236A | S312 | SAMEA104679836 | Italy; Elba; Fetovaia | 42.7313 °N | 10.1534 °E | 2013-07-12 | LPFa |
| MG | 1236B | S313 | SAMEA104679837 | Italy; Elba; Fetovaia | 42.7313 °N | 10.1534 °E | 2013-07-12 | LPFa |
| MG | 1236C | S426 | SAMEA104679838 | Italy; Elba; Fetovaia | 42.7313 °N | 10.1534 °E | 2013-07-19 | LPFa |
| MG | 1418A | M6 | SAMEA104679839 | Italy; Elba; Cavoli | 42.734192 °N | 10.185868 °E | 2014-06-08 | LFY |
| MG | 1418B | M7 | SAMEA104679840 | Italy; Elba; Cavoli | 42.734192 °N | 10.185868 °E | 2014-06-08 | LFY |
| MG | 1418C | BY7 | SAMEA104679841 | Italy; Elba; Cavoli | 42.734192 °N | 10.185868 °E | 2014-11-06 | LFY |
| MG | 1418D | BY1 | SAMEA104679842 | Italy; Elba; Sant' Andrea | 42.808561 °N | 10.142275 °E | 2014-11-03 | TUN |
| MG | 1418E | BY2 | SAMEA104679843 | Italy; Elba; Sant' Andrea | 42.808561 °N | 10.142275 °E | 2014-11-03 | TUN |
| MG | 1418F | BY3 | SAMEA104679844 | Italy; Elba; Sant' Andrea | 42.808561 °N | 10.142275 °E | 2014-11-03 | TUN |
| MG | 1418G | BY8 | SAMEA104679845 | Italy; Elba; Cavoli | 42.734192 °N | 10.185868 °E | 2014-11-06 | UNK |
| MG | 1418H | BY19 | SAMEA104679846 | Italy; Elba; Cavoli | 42.734192 °N | 10.185868 °E | 2014-11-12 | UNK |
| MG | 1743A | BZ163 | SAMEA104679847 | Belize, Twin Cayes, Fisheries Beach | 16.82356 °N | 88.10615 °W | 2015-07-05 | FM |
| MG | 1743B | BZ164 | SAMEA104679848 | Belize, Twin Cayes, Fisheries Beach | 16.82356 °N | 88.10615 °W | 2015-07-05 | FM |
| MG | 1743C | BZ165 | SAMEA104679849 | Belize, Twin Cayes, Fisheries Beach | 16.82356 °N | 88.10615 °W | 2015-07-05 | FM |
| MG | 1743D | BZ157 | SAMEA104679850 | Belize, Twin Cayes, Fisheries Beach | 16.82356 °N | 88.10615 °W | 2015-07-05 | G |
| MG | 1743E | BZ180 | SAMEA104679851 | Belize, Twin Cayes, | 16.82356 | 88.10615 | 2015-07- | G |

| | | | | | | | | |
|---|---|---|---|---|---|---|---|---|
| | | | | Fisheries Beach | °N | °W | 05 | |
| MG | 1743F | BZ181 | SAMEA104679852 | Belize, Twin Cayes, Fisheries Beach | 16.82356 °N | 88.10615 °W | 2015-07-05 | G |
| MG | 1821A | BZ15 | SAMEA104679853 | Belize, Twin Cayes, Fisheries Beach | 16.82383 °N | 88.10582 °W | 2015-06-28 | FW |
| MG | 1821B | BZ106 | SAMEA104679854 | Belize, Twin Cayes, Fisheries Beach | 16.82383 °N | 88.10582 °W | 2015-07-02 | FW |
| MG | 1821C | BZ131 | SAMEA104679855 | Belize, Twin Cayes, Fisheries Beach | 16.82383 °N | 88.10582 °W | 2015-07-03 | FW |
| MG | 1821D | BZ123 | SAMEA104679856 | Belize, Twin Cayes, Fisheries Beach | 16.82383 °N | 88.10582 °W | 2015-07-03 | TC |
| MG | 1821E | BZ125 | SAMEA104679857 | Belize, Twin Cayes, Fisheries Beach | 16.82383 °N | 88.10582 °W | 2015-07-03 | TC |
| MG | 1821F | BZ126 | SAMEA104679858 | Belize, Twin Cayes, Fisheries Beach | 16.82383 °N | 88.10582 °W | 2015-07-03 | TC |
| MG | 1821G | BZ197 | SAMEA104679859 | Belize, Twin Cayes, Fisheries Beach | 16.82356 °N | 88.10615 °W | 2015-07-07 | MB |
| MG | 1821H | BZ198 | SAMEA104679860 | Belize, Twin Cayes, Fisheries Beach | 16.82356 °N | 88.10615 °W | 2015-07-07 | MB |
| MG | 1821I | BZ199 | SAMEA104679861 | Belize, Twin Cayes, Fisheries Beach | 16.82356 °N | 88.10615 °W | 2015-07-07 | MB |
| MG | 2373B | DK47 | SAMEA104693958 | Denmark, Nivå Bay, South of pier | 55.928611 °N | 12.522778 °E | 2016-08-16 | DK |
| MG | 2373C | DK161 | SAMEA104693959 | Denmark, Nivå Bay, South from public beach | 55.936022 °N | 12.525797 °E | 2016-08-18 | DK |
| T | 1056N | SA332 | SAMEA104684497 | Italy; Elba; Sant' Andrea | 42.808561 °N | 10.142275 °E | 2013-08-12 | H |
| T | 1056O | SA331 | SAMEA104684498 | Italy; Elba; Sant' Andrea | 42.808561 °N | 10.142275 °E | 2013-08-12 | H |
| T | 1056P | SA333 | SAMEA104684499 | Italy; Elba; Sant' Andrea | 42.808561 °N | 10.142275 °E | 2013-08-12 | H |
| T | 1056K | S4111 | SAMEA104684500 | Italy; Elba; Fetovaia | 42.7313 °N | 10.1534 °E | 2013-06-18 | SD |
| T | 1056L | S4112 | SAMEA104684501 | Italy; Elba; Fetovaia | 42.7313 °N | 10.1534 °E | 2013-06-18 | SD |
| T | 1056M | S4110 | SAMEA104684502 | Italy; Elba; Fetovaia | 42.7313 °N | 10.1534 °E | 2013-06-18 | SD |

**Supplementary Table 2**. Summary statistics of Kentron genome assemblies. Completeness, contamination, and strain heterogeneity values were estimated with conserved set of marker genes for Gammaproteobacteria using the CheckM pipeline.

| **Library** | **IMG GOLD ID** | **Host sp.** | **Scaff-olds** | **Length (Mbp)** | **GC (%)** | **N50 (kbp)** | **No. ORFs (IMG)** | **Complet eness** | **Conta m-ination** | **Strain hetero-geneity** |
|---|---|---|---|---|---|---|---|---|---|---|
| 1236A | Ga0070988 | LPFa | 1187 | 4.08 | 52.9 | 10.2 | 4823 | 93.68 | 2.86 | 0 |
| 1236B | Ga0070989 | LPFa | 757 | 4.19 | 53.6 | 15.4 | 4389 | 93.54 | 1.31 | 0 |
| 1236C | Ga0070990 | LPFa | 1104 | 4.03 | 52.99 | 10.4 | 4665 | 93.71 | 2.43 | 0 |
| 1418A | Ga0070994 | LFY | 390 | 3.45 | 53.52 | 25.5 | 3533 | 94.37 | 0.89 | 33.33 |
| 1418B | Ga0070995 | LFY | 420 | 3.42 | 53.54 | 26.1 | 3434 | 93.81 | 0.75 | 0 |
| 1418C | Ga0070996 | LFY | 431 | 3.66 | 53.51 | 26.4 | 3768 | 93.84 | 0.75 | 0 |
| 1418D | Ga0071000 | TUN | 909 | 3.62 | 49.78 | 14.4 | 3758 | 94.94 | 1.03 | 0 |
| 1418E | Ga0071001 | TUN | 790 | 3.63 | 49.76 | 17.2 | 3795 | 94.94 | 2.72 | 16.67 |
| 1418F | Ga0071002 | TUN | 772 | 3.62 | 49.76 | 17.1 | 3752 | 94.94 | 2.72 | 16.67 |
| 1418G | Ga0071005 | UNK | 610 | 4.23 | 53.53 | 21.1 | 4318 | 93.93 | 3.28 | 0 |
| 1418H | Ga0071006 | UNK | 609 | 4.22 | 53.6 | 22 | 4325 | 93.93 | 3.46 | 0 |
| 1743A | Ga0114220 | FM | 1923 | 5.01 | 54.17 | 5.9 | 4619 | 93.25 | 3.56 | 37.5 |
| 1743B | Ga0114221 | FM | 1794 | 4.94 | 54.24 | 5.9 | 4560 | 93.81 | 3.56 | 37.5 |
| 1743C | Ga0114222 | FM | 1828 | 5 | 54.16 | 5.9 | 4622 | 93.25 | 3.56 | 37.5 |
| 1743D | Ga0114223 | G | 2741 | 4.75 | 54.16 | 3.7 | 4627 | 94.03 | 3.56 | 14.29 |
| 1743E | Ga0114224 | G | 2723 | 4.65 | 54.21 | 3.7 | 4570 | 94.07 | 3 | 0 |
| 1743F | Ga0114225 | G | 2677 | 4.6 | 54.27 | 3.5 | 4379 | 93.5 | 3 | 0 |
| 1821A | Ga0114235 | FW | 386 | 3.65 | 53.11 | 25.4 | 3533 | 92.97 | 1.31 | 0 |
| 1821B | Ga0114236 | FW | 522 | 3.77 | 53.1 | 23.8 | 3628 | 94.37 | 1.59 | 25 |
| 1821C | Ga0114237 | FW | 454 | 4.05 | 53.42 | 32.3 | 3916 | 94.9 | 1.31 | 0 |
| 1821D | Ga0114238 | TC | 584 | 3.32 | 53.73 | 25.8 | 3425 | 93.91 | 1.31 | 0 |
| 1821E | Ga0114239 | TC | 504 | 3.35 | 53.84 | 36.2 | 3400 | 93.91 | 1.31 | 0 |
| 1821F | Ga0114240 | TC | 558 | 3.31 | 53.74 | 33.5 | 3357 | 93.91 | 1.31 | 0 |
| 1821G | Ga0114241 | MB | 520 | 3.88 | 52.65 | 37.5 | 3630 | 93.22 | 1.87 | 0 |
| 1821H | Ga0114242 | MB | 473 | 3.91 | 52.61 | 36.8 | 3667 | 93.22 | 1.87 | 0 |
| 1821I | Ga0114274 | MB | 467 | 3.93 | 52.56 | 36.8 | 3702 | 93.22 | 1.87 | 0 |
| 2373B | Ga0170837 | DK | 542 | 5.02 | 56.87 | 30.7 | 4506 | 94.11 | 2.25 | 0 |
| 2373C | Ga0170839 | DK | 507 | 4.82 | 56.89 | 35.1 | 4288 | 92.26 | 2.25 | 0 |
| 772A | Ga0070896 | H | 1295 | 3.92 | 55.15 | 10.7 | 4568 | 91.44 | 1.59 | 40 |
| 772B | Ga0070898 | H | 1253 | 3.91 | 55.18 | 10.8 | 4631 | 91.44 | 1.45 | 25 |
| 772C | Ga0070978 | H | 1293 | 3.91 | 55.15 | 10.6 | 4517 | 91.44 | 1.45 | 25 |

| | | | | | | | | | | |
|---|---|---|---|---|---|---|---|---|---|---|
| 772D | Ga0070982 | SD | 579 | 3.81 | 53.45 | 24 | 3963 | 92.46 | 1.31 | 0 |
| 772E | Ga0070983 | SD | 666 | 3.76 | 53.61 | 21.7 | 3913 | 92.46 | 1.87 | 0 |
| 772F | Ga0070984 | SD | 642 | 3.8 | 53.59 | 20.4 | 3934 | 92.46 | 1.87 | 0 |

**Supplementary Table 3.** Key enzymes for autotrophic pathways, and enzymes of reference set used for comparison of read mapping vs SwissProt database. *Sequences for these enzymes were not represented in SwissProt, therefore additional sequences from UniProtKB were added to the database (Supplementary File 4) before mapping.

| Type | Pathway | Enzyme | EC |
|---|---|---|---|
| Autotrophy | Reductive pentose phosphate cycle (Calvin-Benson-Bassham) | Ribulose-1,5-bisphosphate carboxylase/oxidase (RuBisCO) | 4.1.1.39 |
| Autotrophy | Reductive citric acid cycle | ATP citrate lyase (ACL) | (2.3.3.8)* |
| Autotrophy | Reductive citric acid cycle | Citryl-CoA lyase (CCL), Citryl-CoA synthase (CCS) | * |
| Autotrophy | Reductive acetyl-CoA pathway | CO-methylating acetyl-CoA synthase (AcsB) | 2.3.1.169 |
| Autotrophy | 3-Hydroxypropionate bi-cycle | Bifunctional malonyl-CoA reductase (Mcr) | * |
| Autotrophy | 3-Hydroxypropionate bi-cycle | Trifunctional propionyl-CoA synthase (Pcs) | * |
| Autotrophy | 3-Hydroxypropionate/4-hydroxybutyrate cycle | 4-Hydroxybutanoyl-CoA dehydratase (AbfD) | 4.2.1.120 |
| Autotrophy | Dicarboxylate/4-hydroxybutyrate cycle | 4-Hydroxybutanoyl-CoA dehydratase (AbfD) | 4.2.1.120 |
| Reference | Tricarboxylic acid cycle | Malate dehydrogenase | 1.1.1.37 |
| Reference | Tricarboxylic acid cycle | Isocitrate dehydrogenase | 1.1.1.42 |
| Reference | Tricarboxylic acid cycle | 2-Oxoglutarate dehydrogenase | 1.2.4.2 |
| Reference | Tricarboxylic acid cycle | 2-Oxoacid oxidoreductase (ferredoxin) | 1.2.7.11 |
| Reference | Tricarboxylic acid cycle | 2-Oxoglutarate ferredoxin oxidoreductase | 1.2.7.3 |
| Reference | Tricarboxylic acid cycle | Succinate dehydrogenase | 1.3.5.1 |
| Reference | Tricarboxylic acid cycle | Fumarate reducatse (quinol) | 1.3.5.4 |
| Reference | Tricarboxylic acid cycle | Dihydrolipoate dehydrogenase | 1.8.1.4 |
| Reference | Tricarboxylic acid cycle | Dihydrolipoamide S-succinyltransferase | 2.3.1.61 |
| Reference | Tricarboxylic acid cycle | Citrate synthase | 2.3.3.1, 2.3.3.16 |
| Reference | Tricarboxylic acid cycle | Fumarate hydratase | 4.2.1.2 |
| Reference | Tricarboxylic acid cycle | Aconitate hydratase | 4.2.1.3 |
| Reference | Tricarboxylic acid cycle | Succinyl-CoA sytnhetase | 6.2.1.5 |
| Reference | 3-Hydroxypropionate cycle (partial) | Succinyl-CoA-L-malate CoA-transferase | 2.8.3.22 |
| Reference | 3-Hydroxypropionate cycle (partial) | Malyl-CoA lyase | 4.1.3.24 |
| Reference | 3-Hydroxypropionate cycle (partial) | (S)-citramalyl-CoA lyase | 4.1.3.25 |
| Reference | 3-Hydroxypropionate cycle (partial) | Mesaconyl-C1-CoA hydratase | 4.2.1.148 |
| Reference | 3-Hydroxypropionate cycle (partial) | Mesaconyl-C4-CoA hydratase | 4.2.1.153 |
| Reference | 3-Hydroxypropionate cycle (partial) | Mesaconyl-CoA C1-C4 transferase | 5.4.1.3 |
| Reference | 3-Hydroxypropionate cycle (partial) | Methylmalonyl-CoA mutase | 5.4.99.2 |

**Supplementary Table 4**. Potential substrates for Kentron and their oxidation/reduction values.

| Compound | Role | Formula | Oxidation/reduction value |
|---|---|---|---|
| Malic acid | Potential substrate | $C_4H_6O_5$ | +2 |
| Succinic acid | Potential substrate | $C_4H_6O_4$ | +1 |
| Acetic acid | Potential substrate | $C_2H_4O_2$ | 0 |
| Glycogen | Storage compound | $C_{24}H_{42}O_{21}$ | 0 |
| Biomass | | $CH_{1.77}O_{0.49}N_{0.24}$ | -0.395 |
| Propionic acid | Potential substrate | $C_3H_6O_2$ | -1 |
| Polyhydroxybutyrate | Storage compound | $C_4H_6O_2$ | -1 |

**Supplementary Table 5.** List of Transporter Classification families of energy-dependent organic substrate uptake transporters.

| TC | Name |
|---|---|
| 2.A.1.1 | Sugar Porter (SP) Family |
| 2.A.1.4 | Organophosphate:Pi Antiporter (OPA) Family |
| 2.A.1.5 | Oligosaccharide:H+ Symporter (OHS) Family |
| 2.A.1.6 | Metabolite:H+ Symporter (MHS) Family |
| 2.A.1.7 | Fucose: H+ Symporter (FHS) Family |
| 2.A.1.10 | Nucleoside: H+ Symporter (NHS) Family |
| 2.A.1.11 | Oxalate:Formate Antiporter (OFA) Family |
| 2.A.1.12 | Sialate:H+ Symporter (SHS) Family |
| 2.A.1.13 | Monocarboxylate Transporter (MCT) Family (Halestrap, 2011) |
| 2.A.1.14 | Anion:Cation Symporter (ACS) Family |
| 2.A.1.15 | Aromatic Acid:H+ Symporter (AAHS) Family |
| 2.A.1.18 | Polyol Porter (PP) Family |
| 2.A.1.25 | Peptide-Acetyl-Coenzyme A Transporter (PAT) Family |
| 2.A.1.27 | Phenyl Propionate Permease (PPP) Family |
| 2.A.1.52 | Glycerophhosphodiester Uptake (GlpU) Family |
| 2.A.1.53 | Proteobacterial Intraphagosomal Amino Acid Transporter (Pht) Family |
| 2.A.1.56 | 1,3-Dihydroxybenzene Transporter (DHB-T) Family |
| 2.A.1.57 | Ferripyochelin Transporter (FptX) Family |
| 2.A.1.60 | Rhizopine-related MocC (MocC) Family |
| 2.A.1.68 | Glucose Transporter (GT) Family |
| 2.A.2 | Glycoside-Pentoside-Hexuronide (GPH):Cation Symporter Family |
| 2.A.3 | Amino Acid-Polyamine-Organocation (APC) Superfamily |
| 2.A.7.5 | Glucose/Ribose Porter (GRP) Family |
| 2.A.7.6 | L-Rhamnose Transporter (RhaT) Family |
| 2.A.7.18 | Choline Uptake Transporter (LicB-T) Family |
| 2.A.8 | Gluconate:H+ Symporter (GntP) Family |
| 2.A.10 | 2-Keto-3-Deoxygluconate Transporter (KdgT) Family |
| 2.A.11 | Citrate-Mg2+:H+ (CitM) Citrate-Ca2+:H+ (CitH) Symporter (CitMHS) Family |
| 2.A.12 | ATP:ADP Antiporter (AAA) Family |
| 2.A.13 | C4-Dicarboxylate Uptake (Dcu) Family |
| 2.A.14 | Lactate Permease (LctP) Family |
| 2.A.15 | Betaine/Carnitine/Choline Transporter (BCCT) Family |
| 2.A.17 | Proton-dependent Oligopeptide Transporter (POT/PTR) Family |
| 2.A.21 | Solute:Sodium Symporter (SSS) Family |

2.A.22 Neurotransmitter:Sodium Symporter (NSS) Family
2.A.23 Dicarboxylate/Amino Acid:Cation (Na+ or H+) Symporter (DAACS) Family
2.A.24 2-Hydroxycarboxylate Transporter (2-HCT) Family
2.A.25 Alanine or Glycine:Cation Symporter (AGCS) Family
2.A.26 Branched Chain Amino Acid:Cation Symporter (LIVCS) Family
2.A.27 Glutamate:Na+Symporter (ESS) Family
2.A.39 Nucleobase:Cation Symporter-1 (NCS1) Family
2.A.40 Nucleobase/Ascorbate Transporter (NAT) or Nucleobase:Cation Symporter-2 (NCS2) Family
2.A.41 Concentrative Nucleoside Transporter (CNT) Family
2.A.42 Hydroxy/Aromatic Amino Acid Permease (HAAAP) Family
2.A.46 Benzoate:H Symporter (BenE) Family
2.A.47 Divalent Anion:Na+ Symporter (DASS) Family
2.A.50 Glycerol Uptake (GUP) Family
2.A.56 Tripartite ATP-independent Periplasmic Transporter (TRAP-T) Family
2.A.61 C4-dicarboxylate Uptake C (DcuC) Family
2.A.66.2 Polysaccharide Transport (PST) Family
2.A.67 Oligopeptide Transporter (OPT) Family
2.A.68 p-Aminobenzoyl-glutamate Transporter (AbgT) Family
2.A.70 Malonate:Na+ Symporter (MSS) Family
2.A.71 Folate-Biopterin Transporter (FBT) Family
2.A.73 Short Chain Fatty Acid Uptake (AtoE) Family
2.A.80 Tripartite Tricarboxylate Transporter (TTT) Family
2.A.87 Prokaryotic Riboflavin Transporter (P-RFT) Family
2.A.88 Vitamin Uptake Transporter (VUT or ECF) Family
2.A.95 6TMS Neutral Amino Acid Transporter (NAAT) Family
2.A.96 Acetate Uptake Transporter (AceTr) Family
2.A.101 Malonate Uptake (MatC) Family (Formerly UIT1)
2.A.102 4-Toluene Sulfonate Uptake Permease (TSUP) Family
2.A.114 Peptide Transporter Carbon Starvation CstA (CstA) Family
2.A.120 Putative Amino Acid Permease (PAAP) Family
3.A.1.1 Carbohydrate Uptake Transporter-1 (CUT1) Family
3.A.1.2 Carbohydrate Uptake Transporter-2 (CUT2) Family
3.A.1.3 Polar Amino Acid Uptake Transporter (PAAT) Family
3.A.1.4 Hydrophobic Amino Acid Uptake Transporter (HAAT) Family
3.A.1.5 Peptide/Opine/Nickel Uptake Transporter (PepT) Family
3.A.1.11 Polyamine/Opine/Phosphonate Uptake Transporter (POPT) Family
3.A.1.12 Quaternary Amine Uptake Transporter (QAT) Family (Similar to 3.A.1.16 and 3.A.1.17)
3.A.1.13 Vitamin B12 Uptake Transporter (B12T) Family (Similar to 3.A.1.14)

| | |
|---|---|
| 3.A.1.17 | Taurine Uptake Transporter (TauT) Family (Similar to 3.A.1.12 and 3.A.1.16) |
| 3.A.1.19 | Thiamin Uptake Transporter (ThiT) Family (Most similar to 3.A.1.10, 3.A.1.6 and 3.A.1.8 in that order) |
| 3.A.1.20 | Brachyspira Iron Transporter (BIT) Family (Most similar to 3.A.1.6, 3.A.1.8 and 3.A.1.11) |
| 3.A.1.24 | Methionine Uptake Transporter (MUT) Family (Similar to 3.A.1.3 and 3.A.1.12) |
| 3.A.1.25 | Biotin Uptake Transporter (BioMNY) Family |
| 3.A.1.26 | Putative Thiamine Uptake Transporter (ThiW) Family |
| 3.A.1.27 | gamma-Hexachlorocyclohexane (HCH) Family (Similar to 3.A.1.12 and 3.A.1.24) |
| 3.A.1.28 | Queuosine (Queuosine) Family |
| 3.A.1.29 | Methionine Precursor (Met-P) Family |
| 3.A.1.30 | Thiamin Precursor (Thi-P) Family |
| 3.A.1.32 | Cobalamin Precursor (B12-P) Family |
| 3.A.1.33 | Methylthioadenosine (MTA) Family |
| 3.A.1.34 | Tryptophan (TrpXYZ) Family |
| 4.A.1 | PTS Glucose-Glucoside (Glc) Family |
| 4.A.2 | PTS Fructose-Mannitol (Fru) Family |
| 4.A.3 | PTS Lactose-N,N'-Diacetylchitobiose-beta-glucoside (Lac) Family |
| 4.A.4 | PTS Glucitol (Gut) Family |
| 4.A.5 | PTS Galactitol (Gat) Family |
| 4.A.6 | PTS Mannose-Fructose-Sorbose (Man) Family |
| 4.A.7 | PTS L-Ascorbate (L-Asc) Family |
| 4.B.1 | Nicotinamide Ribonucleoside (NR) Uptake Permease (PnuC) Family |
| 4.C.1 | Fatty Acid Group Translocation (FAT) Family |
| 4.C.2 | Carnitine O-Acyl Transferase (CrAT) Family |
| 9.A.5 | Putative Arginine Transporter (ArgW) Family |
| 9.A.18 | Peptide Uptake Permease (PUP) Family |
| 9.A.23 | Niacin/Nicotinamide Transporter (NNT) Family |
| 9.A.28 | Ethanolamine Facilitator (EAF) Family |

**Supplementary Table 6**. Genomes of basal Gammaproteobacteria used for phylogenetic analysis and comparison of organic uptake transporter content. References are to genome description, if published, otherwise to author and date of data deposition. Taxonomy based on LPSN (Euzéby, 1997), if available. Accession numbers are for INSDC contig sets or assemblies unless otherwise indicated.

| **Name** | **Metabolism** | **Reference** | **Taxonomy** | **Accession no.** |
|---|---|---|---|---|
| *Alkalilimnicola ehrlichii* MLHE-1 | Heterotroph | (Hoeft et al., 2007) | Chromatiales; Ectothiorhodospiraceae | GCA_000014785.1 |
| *Allochromatium vinosum* DSM 180 | Photolithotroph | (Weissgerber et al., 2011) | Chromatiales; Chromatiaceae | GCA_000025485.1 |
| *Aquisalimonas asiatica* strain CGMCC 1.6291 | Heterotroph | N. Varghese, 2016 | Chromatiales; Ectothiorhodospiraceae | FOEG01000000 |
| *Arhodomonas aquaeolei* DSM 8974 | Heterotroph | N. Kyrpides et al., 2013 | Chromatiales; Ectothiorhodospiraceae | GCA_000374645.1 |
| *Arsukibacterium* sp. MJ3 | Heterotroph | (Lylloff et al., 2015) | Chromatiales; Chromatiaceae | LAHP01000000 |
| *Beggiatoa alba* B18LD | Chemolithotroph | S. Lucas et al., 2011 | Thiotrichales | GCA_000245015.1 |
| *Beggiatoa* sp. PS | Chemolithotroph | (Mußmann et al., 2007) | Thiotrichales | ABBZ01000000 |
| *Candidatus* Achromatium palustre | Chemolithotroph | (Salman et al., 2016) | Thiotrichales | LFCU01000000 |
| *Candidatus* Competibacter dentrificans | Heterotroph | (McIlroy et al., 2014) | Gammaproteobacteri inc. sed. | CBTJ02000000 |
| *Cardiobacterium hominis* ATCC 15826 | Heterotroph | X. Qin et al. 2009 | Cardiobacterales | GCA_000160655.1 |
| *Coxiella burnetii* RSA 331 | Heterotroph | R. Seshadri, J.E. Samuel, 2007 | Legionellales | GCA_000018745.1 |
| *Cycloclasticus* sp. P1 | Heterotroph | (Lai et al., 2012) | Thiotrichales | GCA_000299965.1 |
| *Dichelobacter nodosus* | Heterotroph | (Myers et al., 2007) | Cardiobacterales | GCA_000015345.1 |
| *Diplorickettsia massiliensis* 20B | Heterotroph | (Mathew et al., 2012) | Legionellales | AJGC01000000 |
| *Ectothiorhodospira haloalkaliphila* ATCC 51935 | Photolithotroph | D.A. Bryant et al., 2013 | Chromatiales; Ectothiorhodospiraceae | AJUE01000000 |

| | | | | |
|---|---|---|---|---|
| *Ectothiorhodosinus mongolicus* strain M9 | Photolithotroph | N. Varghese, 2017 | Chromatiales; Ectothiorhodospiraceae | FTPK01000000 |
| *Ectothiorhodospira* sp. PHS-1 | Photolithotroph | (Kulp et al., 2008) | Chromatiales; Ectothiorhodospiraceae | GCA_000225005.2 |
| *Francisella tularensis* strain FTZ DR87 | Heterotroph | (Davenport et al., 2014) | Thiotrichales | JOVO01000000 |
| *Halorhodospira halophila* SL1 | Photolithotroph | (Challacombe et al., 2013) | Chromatiales; Ectothiorhodospiraceae | GCA_000015585.1 |
| *Halothiobacillus neapolitanus* c2 | Chemolithotroph | | Chromatiales; Halothiobacillaceae | GCA_000024765.1 |
| *Hydrogenovibrio marinus* DSM 11271 | Chemolithotroph | K. Scott et al., 2014 | Thiotrichales | JOML01000000 |
| *Lamprocystis purpurea* DSM 4197 | Photolithotroph | D. Bryant, 2010 | Chromatiales; Chromatiaceae | (JGI/IMG 2515154118) |
| *Leucothrix mucor* DSM 2157 | Chemolithotroph | (Wu et al., 2009) | Thiotrichales | ATTE01000000 |
| *Marichromatium purpuratum* 987 | Photolithotroph | | Chromatiales; Chromatiaceae | (JGI/IMG 2510065050) |
| *Methylococcus capsulatus* str. Bath | Heterotroph | (Ward et al., 2004) | Methylococcales | GCA_000008325.1 |
| *Methylomonas methanica* | Heterotroph | (Boden et al., 2011a) | Methylococcales | GCA_000214665.1 (JGI/IMG 2504756059) |
| *Methylophaga thiooxydans* DMS010 | Chemolithotroph | (Boden et al., 2011b) | Thiotrichales | ABXT01000000 |
| *Methylosarcina fibrata* AML-C10 | Heterotroph | M.G. Kalyuzhnaya et al., 2013 | Methylococcales | GCA_000372865.1 |
| *Nitrococcus mobilis* Nb-231 | Chemolithotroph | J. Waterbury et al., 2006 | Chromatiales; Ectothiorhodospiraceae | AAOF01000000 |
| *Nitrosococcus oceani* ATCC 19707 | Chemolithotroph | (Klotz et al., 2006) | Chromatiales; Chromatiaceae | GCA_000012805.1 |
| Pinkberry Chromatiaceae | Photolithotroph | (Wilbanks et al., 2014) | Chromatiales | (de novo reassembled) |
| *Piscirickettsia salmonis* strain AY3800B | Heterotroph | (Bohle et al., 2017) | Thiotrichales | GCA_001746795.1 |
| *Rheinheimera* sp. A13L | Heterotroph | (Gupta et al., 2011) | Chromatiales; Chromatiaceae | AFHI01000000 |
| *Rheinheimera baltica* DSM 14885 | Heterotroph | N. Kyrpides et al., 2013 | Chromatiales; Chromatiaceae | AUDG01000000 |
| *Rheinheimera nanhaiensis* E407-8 | Heterotroph | (Zhang et al., 2012) | Chromatiales; Chromatiaceae | BAFK01000000 |
| *Spiribacter salinus* M19-40 | Heterotroph | (León et al., 2014) | Chromatiales; Ectothiorhodospiraceae | GCA_000319575.2 |

| | | | | |
|---|---|---|---|---|
| *Sulfurivirga caldicuralii* strain DSM 17737 | Chemolithotroph | N. Varghese, 2016 | Thiotrichales | FSRE01000000 |
| *Thioalkalimicrobium* sp. ALE5 | Chemolithotroph | N. Varghese, 2016 | Thiotrichales | FOYY01000000 |
| *Thioalkalivibrio nitratireducens* DSM 14787 | Chemolithotroph | T.V. Tikhonova et al., 2015 | Chromatiales; Ectothiorhodospiraceae | GCA_000321415.2 |
| *Thiocapsa marina* 5811 | Photolithotroph | S. Lucas et al., 2011 | Chromatiales; Chromatiaceae | GCA_000223985.2 |
| *Thiocystis violascens* DSM 198 | Photolithotroph | S. Lucas et al., 2012 | Chromatiales; Chromatiaceae | GCA_000227745.3 |
| *Thioflavicoccus mobilis* 8321 | Photolithotroph | D. Bryant, 2010 | Chromatiales; Chromatiaceae | (JGI/IMG 2506783059) |
| *Thiohalomonas denitrificans* strain HLD2 | Chemolithotroph | N. Varghese, 2016 | Chromatiales; Ectothiorhodospiraceae | FMWD01000000 |
| *Thiomargarita nelsonii* | Chemolithotroph | (Winkel et al., 2016) | Thiotrichales | JSZA01000000 |
| *Thiomicrospira kuenenii* DSM 12350 | Chemolithotroph | | Thiotrichales | JAGP01000000 |
| *Thioploca ingrica* | Chemolithotroph | (Kojima et al., 2015) | Thiotrichales | GCA_000828835.1 |
| *Thiorhodococcus drewsii* AZ1 | Photolithotroph | S. Lucas et al., 2011 | Chromatiales; Chromatiaceae | AFWT01000000 |
| *Thiorhodospira sibirica* ATCC 700588 | Photolithotroph | S. Lucas et al., 2011 | Chromatiales; Ectothiorhodospiraceae | AGFD01000000 |
| *Thiorhodovibrio* sp. 907 | Photolithotroph | S. Lucas et al., 2011 | Chromatiales; Chromatiaceae | AFWS02000000 |
| *Thiothrix nivea* DSM 5205 | Chemolithotroph | (Lapidus et al., 2011) | Thiotrichales | AJUL01000000 |
| *Wenzhouxiangella marina* strain KCTC 42284 | Heterotroph | (Lee et al., 2015) | Chromatiales; Wenzhouxiangellaceae | GCA_001187785.1 |
| *Xanthomonas oryzae pv. oryzae* MAFF 311018 | Heterotroph | (Ochiai et al., 2005) | Xanthomonadales | GCA_000010025.1 |
| *Xylella fastidiosa subsp. fastidiosa* GB514 | Heterotroph | (Schreiber et al., 2010) | Xanthomonadales | GCA_000148405.1 |
| *Candidatus* Ruthia magnifica str. Cm | Symbiont | (Roeselers et al., 2010; Newton et al., 2007) | Host: *Calyptogena magnifica* (bivalve) | GCA_000015105.1 |
| *Candidatus* Thiodiazotropha endoloripes | Symbiont | (Petersen et al., 2016) | Host: *Loripes lucinalis* (bivalve) | LVJW00000000 |

| | | | | |
|---|---|---|---|---|
| *Candidatus* Thiodiazotropha endolucina | Symbiont | (König et al., 2016) | Host: *Codakia orbicularis* (bivalve) | MARB01000000 |
| *Candidatus* Thiosymbion oneisti | Symbiont | (Petersen et al., 2016) | Host: *Laxus oneistus* (nematode) | FLUY00000000 |
| *Candidatus* Vesicomyosocius okutanii HA | Symbiont | (Kuwahara et al., 2007) | Host: *Calyptogena okutanii* (bivalve) | GCA_000010405.1 |
| Endosymbiont Gamma3 of *Olavius algarvensis* | Symbiont | (Woyke et al., 2006; Kleiner et al., 2011) | Host: *Olavius algarvensis* (oligochaete) | (JGI/IMG 3300005391) |
| Endosymbiont of *Bathymodiolus azoricus* | Symbiont | (Sayavedra et al., 2015) | Host: *Bathymodiolus azoricus* (bivalve) | CDSC02000000 |
| Endosymbiont of *Bathymodiolus* sp. | Symbiont | (Sayavedra et al., 2015; Petersen et al., 2011) | Host: *Bathymodiolus* sp. (bivalve) | CAEB01000000 |
| Endosymbiont of *Riftia pachyptila* | Symbiont | (Robidart et al., 2008) | Host: *Riftia pachyptila* (vent Ph05) (vestimentiferan tubeworm) | AFOC01000000 |
| Endosymbiont of *Solemya velum* strain WH | Symbiont | (Dmytrenko et al., 2014) | Host: *Solemya velum* (bivalve) | JRAA01000000 |
| Endosymbiont of *Tevnia jerichonana* | Symbiont | (Gardebrecht et al., 2012) | Host: *Tevnia jerichonana* (vent Tica) (vestimentiferan tubeworm) | AFZB01000000 |
| Endosymbiont of unidentified scaly snail | Symbiont | (Nakagawa et al., 2014) | Host: Unidentified scaly snail isolate Monju (cf. *Chrysomallon*) | GCA_000801295.1 |

**Supplementary Table 7**. Collection localities and dates for *Kentrophoros* metabolomics samples.

| Sample no. | No. cells | Locality | Latitude | Longitude | Date | Host phylotype |
|---|---|---|---|---|---|---|
| L2 | 5 | Italy; Elba; Cavoli | 42.734192 °N | 10.185868 °E | 2014-11-06 | H |
| L3 | 10 | Italy; Elba; Cavoli | 42.734192 °N | 10.185868 °E | 2014-11-06 | H |
| L4 | 10 | Italy; Elba; Cavoli | 42.734192 °N | 10.185868 °E | 2014-11-07 | H |
| L12 | 12 | Italy; Elba; Cavoli | 42.734192 °N | 10.185868 °E | 2014-11-12 | H |
| L13 | 10 | Italy; Elba; Cavoli | 42.734192 °N | 10.185868 °E | 2014-11-12 | H |

**Supplementary Table 8**. Hypothetical reaction scheme that can allow autotrophic $CO_2$ fixation with enzymes that are predicted in Kentron genomes. Free energy values ($\Delta_r G'^m$) were calculated for pH 7.0 and concentrations 1 mM using eQuilibrator(17).

| **Enzyme** | **EC** | **Reaction** | **Flux** | **$\Delta_r G'^m$** (kJ/mol) | **+/-** |
|---|---|---|---|---|---|
| MMC lyase | 4.1.3.24 | (S)-Malyl-CoA ⇌ Glyoxylate + Acetyl-CoA | 1 | -4.2 | 5.8 |
| MMC lyase | 4.1.3.24 | Glyoxylate + Propionyl-CoA ⇌ Beta-Methylmalyl-CoA | 1 | 2.8 | 16.4 |
| Mesaconyl-C1-CoA hydratase (Mch) | 4.2.1.148 | Beta-Methylmalyl-CoA ⇌ Mesaconyl-C1-CoA + H2O | 1 | -3.2 | 3.7 |
| Mesaconyl-CoA-C1:C4 CoA transferase (Mct) | 5.4.1.3 | Mesaconyl-C1-CoA ⇌ Mesaconyl-C4-CoA | 1 | 0.0 | 0.0 |
| Mesaconyl-C4-CoA hydratase (Meh) | 4.2.1.153 | Mesaconyl-C4-CoA + H2O ⇌ (S)-Citramalyl-CoA | 1 | -3.2 | 3.7 |
| MMC lyase | 4.1.3.25 | (3S)-Citramalyl-CoA ⇌ Acetyl-CoA + Pyruvate | 1 | -7.9 | 15.3 |
| Pyruvate synthase (PFOR) | 1.2.7.1 | Acetyl-CoA + **$CO_2$** + 2 Ferredoxin (red) ⇌ **Pyruvate** + 2 Ferredoxin (ox) + CoA | 2 | 18.7 | 13.2 |
| Pyruvate phosphate dikinase (PPDK) | 2.7.9.1 | Pyruvate + ATP + Orthophosphate ⇌ Phosphoenolpyruvate + AMP + Pyrophosphate | 1 | 19.6 | 1.0 |
| PEP carboxykinase (GDP) | 4.1.1.32 | Phosphoenolpyruvate + GDP + **$CO_2$** ⇌ Oxaloacetate + GTP | 1 | 4.3 | 6.7 |
| Methylmalonyl-CoA carboxytransferase (MMCT) | 2.1.3.1 | Pyruvate + (S)-Methylmalonyl-CoA ⇌ Oxaloacetate + Propionyl-CoA | 1 | -2.3 | 10.4 |
| Malate dehydrogenase | 1.1.1.37 | Oxaloacetate + NADH ⇌ (S)-Malate + $NAD^+$ | 2 | -30.3 | 0.6 |
| Fumarate hydratase (FumH) | 4.2.1.2 | (S)-Malate ⇌ Fumarate + $H_2O$ | 1 | 3.5 | 0.6 |
| Fumarate reductase (FumR) | 1.3.99.1 | Fumarate + $FADH_2$ ⇌ Succinate + FAD | 1 | -46.2 | 7.1 |
| Succinyl-CoA synthase (SCS) | 6.2.1.5 | Succinate + ATP + CoA ⇌ Succinyl-CoA + ADP + Orthophosphate | 2 | -1.8 | 2.6 |
| Methylmalonyl-CoA mutase (MMCM) | 5.4.99.2 | Succinyl-CoA ⇌ (R)-Methylmalonyl-CoA | 1 | 7.6 | 4.1 |
| Methylmalonyl-CoA epimerase (EPI) | 5.1.99.1 | (R)-Methylmalonyl-CoA ⇌ (S)-Methylmalonyl-CoA | 1 | 0.0 | 5.8 |
| Succinyl-CoA:Malate CoA transferase (Smt) | 2.8.3.22 | Succinyl-CoA + (S)-Malate ⇌ Succinate + (S)-Malyl-CoA | 1 | -4.8 | 7.4 |
| | **TOTAL** | 3 $CO_2$ + 4 Ferredoxin (red) + 2 NADH + $FADH_2$ + 3 ATP + GDP ⇌ Pyruvate + 4 Ferredoxin (ox) + 2 $NAD^+$ + FAD + $H_2O$ + 2 ADP + AMP + GTP + Pyrophosphate + Orthophosphate | | -60.9 | 30.0 |

**Supplementary Table 9**. Effect of changing the ratio of reduced:oxidized ferredoxin on the free energy yield for the reductive pyruvate synthase reaction.

| Ferredoxin(red) (mM) | Ferredoxin(ox) (mM) | Ratio red:ox | $\Delta_r G'$ of pyruvate synthase reaction |
|---|---|---|---|
| 1 | 1 | 1 | 18.7 |
| 1 | 0.1 | 10 | 7.3 |
| 1 | 0.01 | 100 | -4.1 |
| 1 | 0.001 | 1000 | -15.5 |

**Supplementary Table 10.** Number of genomes with each predicted metabolism in the IMG/ ER database, based on the presence/absence of key genes (under “Criteria”). Except where noted, the number of genomes was not filtered for genome completeness. * Excluding Kentron and incomplete genomes.

| **Predicted metabolism** | | **Criteria** | | | | **No. genomes** |
|---|---|---|---|---|---|---|
| | | **RbcL** | **Acl or Ccl** | **rDsrAB** | **SoxB,XA,YZ** | |
| Thioheterotroph | rDsr-Sox | - | - | + | + | 7* |
| | Sox | - | - | - | + | 661 |
| Thioautotroph | CBB, rDsr-Sox | + | - | + | + | 89 |
| | CBB, Sox | + | - | - | + | 600 |
| | rTCA, Sox | - | + | - | + | 41 |
| | CBB+rTCA, Sox | + | + | - | + | 1 |
| | CBB + rTCA, rDsr-Sox | + | + | + | + | 8 |

**Supplementary Table 11**. Direct protein stable isotope fingerprinting (SIF) values for Kentron sp. H. $\delta^{13}C$ values were offset-corrected using a human hair standard for each instrument run. * samples excluded because insufficient peptides were detected. Full raw and processed data are available from the PRIDE repository (see *Data availability* in main text).

| **Sample** | **Locality** | **No. host cells** | **$\delta^{13}C$ (‰)** | **Standard error of $\delta^{13}C$ (‰)** | **Peptides passing final filter** | **Unique peptides** |
|---|---|---|---|---|---|---|
| Ind2 | Elba | 1 | -2.5 | 4.7 | 14 | 10 |
| Pool1 | Elba | 5 | -3.1 | 1.9 | 178 | 76 |
| Pool2 | Elba | 5 | -12.3 | 2.3 | 456 | 165 |
| Pool3 | Elba | 4 | -5.25 | 1.3 | 186 | 57 |
| Pool4 | Elba | 5 | -4.75 | 2.4 | 50 | 21 |
| Pool5 | Elba | 5 | -5.05 | 1.5 | 499 | 132 |
| Pool6 * | Elba | 5 | 6.35 * | 12.1 * | 7 | 6 |
| Pool7 * | France | 5 | * | * | 0 | 0 |
| Pool8 | France | 4 | -2.85 | 2.6 | 85 | 32 |
| Pool9 | Elba | 5 | -5.65 | 1.4 | 201 | 54 |

**Supplementary Table 12**. Stable isotope ratios for dissolved inorganic carbon in Elba seawater and sediment porewater.

| **Sample** | **Type** | **$\delta^{13}C$ (‰)** | **Standard error of $\delta^{13}C$ (‰)** |
|---|---|---|---|
| 101 | porewater | -2.42 | 0.06 |
| 102 | porewater | -1.69 | 0.05 |
| 103 | porewater | -1.32 | 0.07 |
| 104 | porewater | -2.99 | 0.05 |
| 105 | porewater | -1.93 | 0.05 |
| 106 | seawater | 0.92 | 0.07 |
| 107 | seawater | 1.55 | 0.03 |
| 108 | seawater | 0.83 | 0.08 |

#### List of Supplementary Files

The following files are available online at: https://doi.org/10.5281/zenodo.2555833

**Supplementary File 1.** Hypothetical reaction scheme for autotrophic $CO_2$ fixation. IMG locus IDs for identified genes, corresponding mean expression levels in Kentron spp. H and SD, and their occurrence in other genomes in IMG/ER database.

**Supplementary File 2.** Transcriptome expression data for Kentron spp. H and SD.

**Supplementary File 3.** Counts of uptake transporter families (by Transporter Classification number) per genome for Kentron and Gammaproteobacteria used for comparison.

**Supplementary File 4.** UniProt accession numbers of key autotrophy enzymes added to SwissProt database.
